# Genes involved in protein folding and chromatin organization buffer genetic variation

**DOI:** 10.1101/2024.09.24.614041

**Authors:** Jens Frickel, Mohammed T. Tawfeeq, Emily Baker, Sara Baco, Jonas Rombout, Karin Voordeckers, Daniel F. Jarosz, Sibylle C. Vonesch, Kevin J. Verstrepen

## Abstract

Mutations are not always phenotypically active or show different effects in different individuals. While the mechanisms underlying this variable relationship between mutations and phenotypes are largely elusive, some specific genes may influence the phenotypic effects of cryptic variation. We employ the toolbox of *Saccharomyces cerevisiae* to perform a genome-wide screen aimed at identifying these so-called genetic buffer genes. Measuring the fitness of 1.8 million mutated strains identified a small set of evolutionary conserved buffer genes involved in protein folding and chromatin organization, including *GIM3, SSA2, HOG1* and *FKH2*. Deletion of these genes increases the fitness effect of de novo mutations as well as standing genetic variation, with some mutations becoming adaptive. Moreover, losing a buffer gene results in a decline of standing genetic variation. Together, these results reveal a set of conserved genes that influence the phenotypic outcome of mutations and play a role in complex genetics and incomplete penetrance.

## INTRODUCTION

Genetic variation fuels organisms’ capacity to adapt to new niches and environmental challenges. While adaptation often requires *de novo* mutations, standing genetic variation, i.e. variation that is already present within a population at a given time, is considered as one of the major factors contributing to adaptation (Barrett and Schluter, 2008). Interestingly, a large part of genetic variation is phenotypically neutral and does not directly affect an organism’s fitness (Gibson and Dworkin, 2004a; Hietpas et al., 2011; Waddington, 1942). However, when the environment changes, some neutral variation can become phenotypically active and contribute to adaptation (Berger et al., 2011; Dworkin et al., 2003; Gibson and Dworkin, 2004b; Lauter and Doebley, 2002; McGuigan et al., 2011; Paaby and Rockman, 2014). One of the best-known experiments that showed the importance of such so-called cryptic genetic variation (CGV) has been Luria and Delbruck’s fluctuation assay. Their research demonstrated that large populations of *Escherichia coli* cells often contain cryptic genetic variants that become adaptive when the cells are exposed to phage T1 because they limit viral docking (Lurias and Delbrock, 1943). Over the years, the list of examples of the importance of CGV has become seemingly endless, with examples ranging from development (Gibson and Hogness, 1996; Waddington, 1953) to the potential of standing CGV as an accelerator of RNA enzyme evolution (Hayden et al., 2011).

Importantly, CGV can not only become phenotypically active upon environmental change, but also because of physiological or genetic changes occurring within the organism. Such genetic interactions help explain why novel mutations can be cryptic in some individuals, but not in others, depending on the genotype of the individual. This so-called incomplete penetrance is for example observed in the context of cancer development and immunodeficiency diseases, where different outcomes of specific mutations have been at least partly attributed to the genetic background in which they occur (Gibson and Dworkin, 2004b; Gruber and Bogunovic, 2020).

Many examples of the interaction between CGV and other mutations or perturbations involve specific and complex combinations of mutations. However, Rutherford and Lindquist’s (Rutherford and Lindquist, 1998) seminal work has revealed that the activity of one specific gene, *HSP90*, can influence the effect of a large number of cryptic mutations. Specifically, reduced activity of the Hsp90 chaperone, either due to drug-based inhibition or natural stochastic fluctuations, results in the appearance of a wide range of heritable phenotypes in organisms ranging from flies to plants due to the exposure of previously hidden CGV (Burga et al., 2011; Casanueva et al., 2012; Chen and Wagner, 2012; Cowen and Lindquist, 2005; Jarosz and Lindquist, 2010; Koneru et al., 2021; Queitsch et al., 2002; Yeyati et al., 2007). *HSP90* is sometimes referred to as a “genetic buffer” due to its potential to hide or buffer the effects of many mutations. It is often hypothesized that the influence of Hsp90 on the phenotypic outcome of CGV may also affect evolutionary adaptation, and several studies have demonstrated a link between Hsp90 activity and the adaptive potential of CGV in natural populations (Casanueva et al., 2012; Paaby and Rockman, 2014; Rohner et al., 2013; Rutherford, 2003; Sangster et al., 2008). Furthermore, complex human diseases have been linked to CGV (Gibson and Dworkin, 2004a; Paaby and Rockman, 2014), and natural fluctuations in the activity of the buffer gene *HSP90* have been linked to disease severity (Hummel et al., 2017; Karras et al., 2017; Siegal, 2017).

Despite the large number of studies that have demonstrated the buffering activity of Hsp90, several seminal questions remain. For example, whether genetic buffers such as *HSP90*, suppress the effect of a broad range of mutations, representing a system level property has remained debated. It has been suggested that buffered cryptic variants might be intrinsically rare and only represent specific cases that gradually accumulate in genomes through purifying selection (Geiler-Samerotte et al., 2016; Richardson et al., 2013; Siegal and Leu, 2014). Moreover, it also remains unclear to what extent *HSP90* is a special, rare example of a buffer gene. Some studies have shown that overexpression of other molecular chaperones can buffer the negative fitness effects associated with high mutational loads (Aguilar-Rodríguez et al., 2016; Fares et al., 2002; Maisnier-Patin et al., 2005; Tokuriki and Tawfik, 2009), suggesting that Hsp90 might not be unique. This indicates a need to understand the prevalence of genetic buffers and their effect on CGV. Research to date has not yet offered a systematic study investigating the prevalence of buffer genes and their influence on genetic variation.

Here, we performed a genome-wide screen for genes that act as genetic buffers, by combining the genetic toolbox of the model eukaryote *Saccharomyces cerevisiae* with high-throughput phenotyping. We identify a large set of candidate buffer genes and show that these are often involved in protein folding and cellular regulation through chromatin modification. Interestingly, our results reveal that impairing the function of these genes not only increases the fitness effects of *de novo* mutations, but also reduces how much genetic variation is retained. These results shed light on the general characteristics of genetic buffers, and show that these buffers influence both the maintenance as well as the de-silencing of genetic variation. Moreover, the buffering genes that are identified in this study are often evolutionary conserved and might thus also play a role in incomplete penetrance and complex genetic interactions across the tree of life.

## RESULTS

We combined high-throughput robotics, automated single-cell microscopy and image analysis with the molecular toolbox for the model eukaryote *S. cerevisiae*. The aim was development of a genome-wide screen to identify genes that act as a genetic buffer. We measured the fitness effect of random mutations in the absence of non-essential genes (Figure 1A) using the haploid yeast gene knockout collection (YKO) combined with mutagenesis. Each of the ∼4500 single-gene deletion strains was cultured and split into a control and mutant culture, which was mutagenized by exposure to UV-light (see Methods) resulting in multiple different random mutations in each cell (Figure 1A and Figure S1). Based on pooled sequencing (Figure S1), cells had on average ∼4 mutations spanning the intergenic and coding regions, resulting both in synonymous and non-synonymous mutations. We reasoned that random *de novo* mutations will frequently evoke a stronger effect on a cell’s fitness in the absence of a buffering gene (Figure 1B). As a proxy for fitness, we used automated microscopy to track the growth of ∼200 individual cells (lineages), for each of the 4500 mutagenized and 4500 un-mutagenized cultures. Based on their growth curves, we calculated the fitness of each lineage (Figure 1A). We used relative fitness (average fitness of each mutagenized culture relative to the average fitness of the corresponding un-mutagenized control culture) as a measure for the buffering potential of a gene, with lower relative fitness of a mutagenized deletion strain indicating a stronger buffering potential for the specific gene that was deleted. We confirmed that these fitness measurements were highly reproducible, because fitness obtained for lineages in control cultures with overlapping ORFs deleted were correlated (Linear model *F*_(1,222)_=558.1, *P* < 2.2e-16, *R^2^*=0.71), This correlation was expected since these two lineages essentially miss the same gene function. Moreover, it was found that the buffering potential of pseudo-genes (genomic regions not coding for a protein) that do not overlap with functional genes was low (‘deletion pseudo-genes’; Figure 2A).

**Figure 1.**
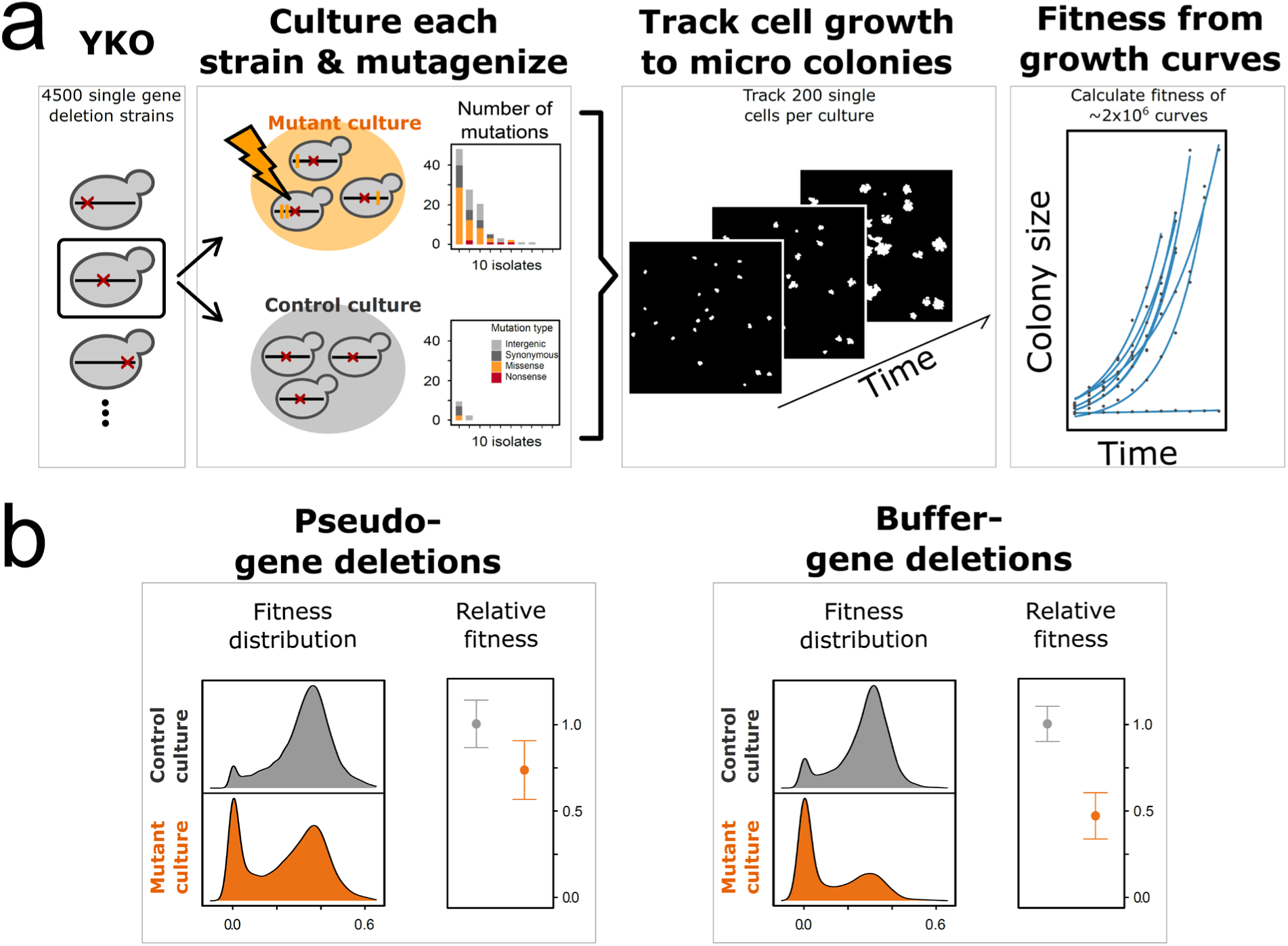
Genome wide screen to identify buffer genes. (A) Schematic representation of the genome wide screen. Around 4500 different strains of the haploid yeast single-gene knockout collection (YKO) were cultured and subsequently split into a ‘control culture’ and ‘mutant culture’. Random mutations were induced in each of the 4500 mutant cultures by exposure to UV light. To estimate the average mutational load in the UV-mutagenized cultures, ten randomly selected isolates from a mutant culture and control culture were sequenced to identify mutations and mutation type in each isolate. As expected, isolates from the mutant culture showed an increased mutational load, and the exact mutations differed between each of the sequenced isolates (see also Figure S1). For each culture, automated microscopy was used to track the growth of around 200 individual single cells. Each hour, pictures were taken as microcolonies were developing, which were processed to growth curves to estimate the fitness of each mutant and un-mutagenized lineage, resulting in around 2,00,000 growth curves. (B) The fitness distribution of random mutations is different when buffer-genes (candidate buffer genes cf. figure 2) are deleted, compared to the deletion of pseudo-genes. The mutant fitness distribution shows a strong increase of cells with low fitness compared those with pseudo-gene deletions. As expected, the average fitness of mutagenized strains is lower compared to the un-mutagenized parent (relative fitness: fitness is relative to the average fitness of un-mutagenized control culture with the same pseudo-gene or buffer gene deleted, error bars represent average standard deviation in relative fitness).On average, random mutations cause a stronger decrease in relative fitness when a buffer gene is deleted compared to the effect when a pseudo-gene is deleted.

**Figure 2.**
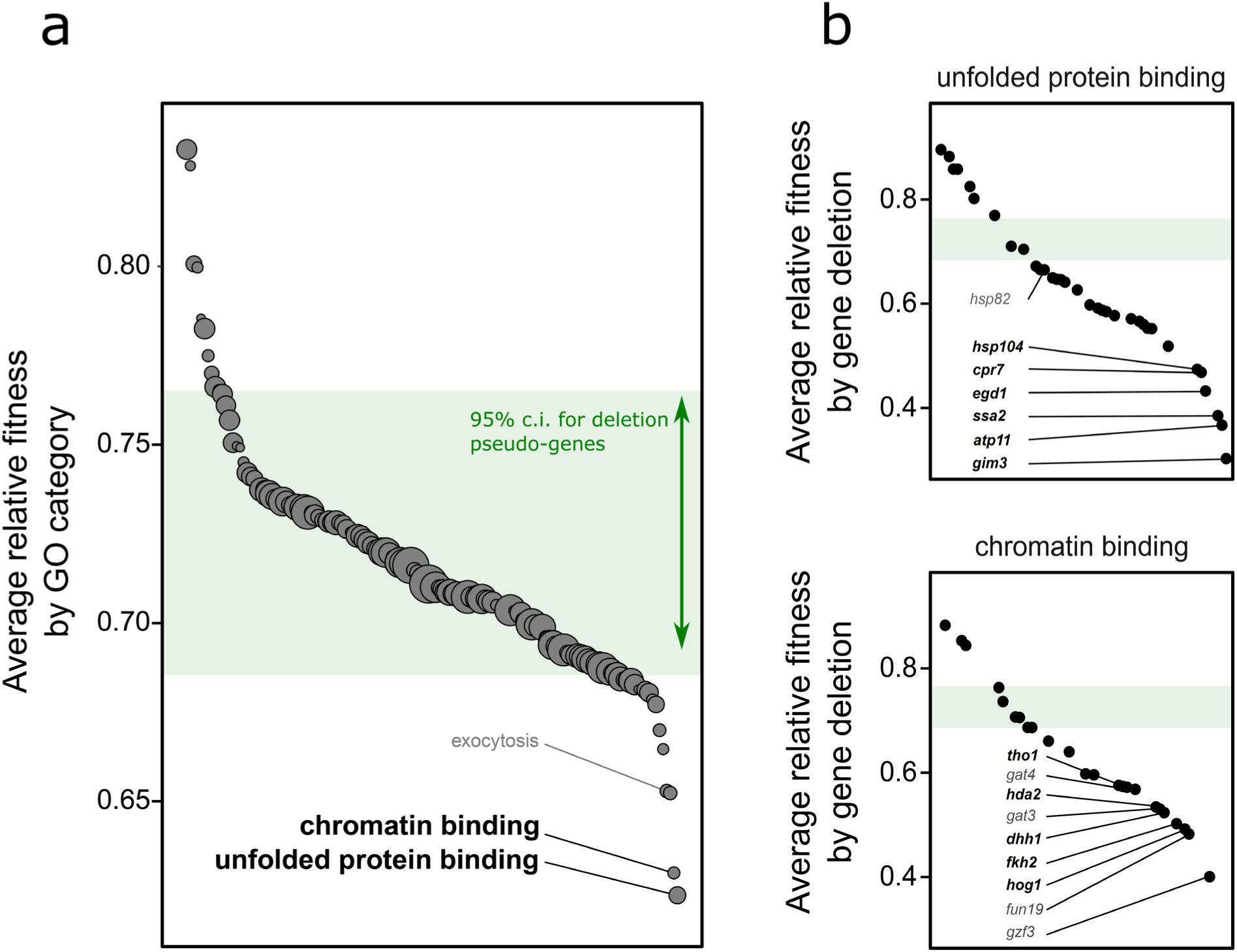
Identification of functional groups and genes with high buffering potential. (A) The buffering capacity of genes grouped by their GO annotations. Note that a lower relative fitness indicates a stronger buffering capacity. Perturbations (by gene deletions) in most functional groups (cf. GO annotations) do not lead to a stronger effect of random mutations compared to the deletion of pseudogenes (green marked area; +/- 95% ci). The functional groups ‘unfolded protein binding’ and ‘chromatin binding’ show a remarkable strong buffering potential. When these molecular functions are perturbed, the average effect of random mutations is much more severe compared to other functional groups or the deletion of pseudo-genes. The size of the dots corresponds to the number of genes contained within each functional group (minimum 20 genes per group). (B) The buffering capacity of individual genes within the two functional groups with strongest buffering potential. Candidate genes for further testing are indicated in bold. Genes indicated in grey were annotated using computational approaches without experimental validation and not selected for further testing.

### Genome-wide screen reveals buffer genes involved in unfolded protein response and chromatin binding

First, we asked in general whether there was a relationship between our measure of buffering potential and known genetic interaction properties of all the tested genes, which are archived in the BioGrid database (Stark et al., 2006). Genes with strong buffering potential had significantly more dosage rescue interactions and synthetic growth defect interactions (GLM, family = Gamma, link = log, *F_(4,3469)_* = 7.0434, *P* = 1.208e-05; dosage rescue: *F_(1,3470)_* = 7.05, *P* = 0.007944581; synthetic growth defect *F_(1,_*_3471)_ = 20.56, *P* = 5.97e-6; See Methods & Table S1). These findings are consistent with expected properties of a genetic buffer, showing that they can mitigate the effect of random mutations.

The next objective was to determine which molecular mechanisms could work as a genetic buffer by grouping all deletion strains (except deletion strains where a DNA-repair gene was deleted, see Methods) according to the GO (gene ontology) annotations of the deleted genes. Most GO categories did not show a greater buffering potential compared to the deletion of pseudo-genes (Figure 2A). There are, however, two functional categories that showed an exceptionally strong buffering potential (i.e. the two GO categories with lowest average relative fitness). The first category consists of genes involved in ‘unfolded protein binding’. This category contains, amongst others, chaperone genes like *HSP90*, previously referred to as a genetic buffer. The second GO category, ‘chromatin binding’ is also of particular interest because it contains genes that are involved in cellular regulation, often through chromatin organization and modification. A previous study found that chromatin regulators can buffer the variation in gene expression between two closely related yeast species, suggesting that they buffer the effect of genetic variation between the two species (Tirosh et al., 2010). Furthermore, we used a GO enrichment analysis to investigate all tested genes ranked by their buffering potential and found a significant enrichment for these two GO terms (amongst others, Table S2: unfolded protein binding, adjusted *P* = 0.008; chromatin binding, adjusted *P* = 0.048).

Interestingly, genes within these two GO categories are relatively highly connected within the genetic network. The average number of genetic interactors was higher compared to all other genes (GLM, family = quasipoisson, link = log, *F_(1,3242)_* = 6.65, *P* = 0.009942). Previous research has shown that loss of highly connected ‘hub’ genes increases environmental variation in genetically identical yeast (Levy et al., 2008). A study that quantified genetic interactions in *C. elegans* found that inactivation of hub genes can enhance phenotypic consequences of mutations in other unrelated genes (Lehner et al., 2006). Many of such genes were chromatin regulators, suggesting that these genes can function as general buffers of both environmental and genetic variation.

### Experimental validation of the buffering function of selected candidate genes

It is noteworthy that not all genes in these two GO categories showed equally strong buffering potential (Figure 2B). For example, although the previously described genetic buffer; *HSP90* (*HSP82* in yeast), had a stronger buffering potential compared to pseudo-genes, other genes within this GO category showed much stronger effects. A potential explanation for this is that the effect of deleting *HSP82* can be (partially) compensated by its paralog *HSC82*, which is constitutively expressed at higher levels. However, since the double mutant is inviable, we could not verify whether this is indeed the case.

To experimentally validate the buffering function, we focused on 11 well-characterized candidate buffer genes involved in ‘unfolded protein binding’ and ‘chromatin binding’ (Figure 2B in bold), by selecting genes with strongest buffering potential. First, we confirmed that deleting these genes neither increased mutation rate nor resulted in increased UV sensitivity (Table S1 & Figure S2); since such strains would acquire more mutations which is not our definition of a buffering gene. Next, two follow-up experiments were used to verify the buffering potential of the selected candidate genes. We isolated ten randomly mutagenized WT (wild type) lineages, each carrying a different set of random mutations (confirmed by sequencing, Figure S1A). Each of the 11 candidate genes were deleted in each of these 10 mutant strains, allowing us to calculate epistasis between mutations and the candidate genes (c.f. Wagih *et al*.(Wagih et al., 2013)). The results revealed several cases of negative epistasis (Figure 3; negative genetic interaction) between the random mutations and our candidate genes, demonstrating that deleting the candidate buffering gene enhanced the (mostly negative) fitness effects of the different sets of random mutations. Furthermore, it was not possible to generate several deletions (Figure 3; red dots). This suggests a synthetic lethal interaction between mutations and the candidate gene, which can be considered as an extreme case of negative epistasis. We also identified a few cases of positive interactions (Figure 3; positive genetic interaction), where deleting a buffer gene in a specific mutagenized strain resulted in a higher-than-expected fitness. However, these positive interactions were much less frequent and less pronounced. Importantly, the set of mutant strains showing negative epistasis differed between the candidate genes, suggesting that the candidate genes do not buffer the same set of mutations. Two candidate genes (*HSP104* and *HDA2*) did not show significant epistasis with any of the tested mutations. These genes had, within their respective GO categories, lower buffering potential compared to the other candidate genes as determined by our genome-wide screen (Figure 2B).

**Figure 3.**
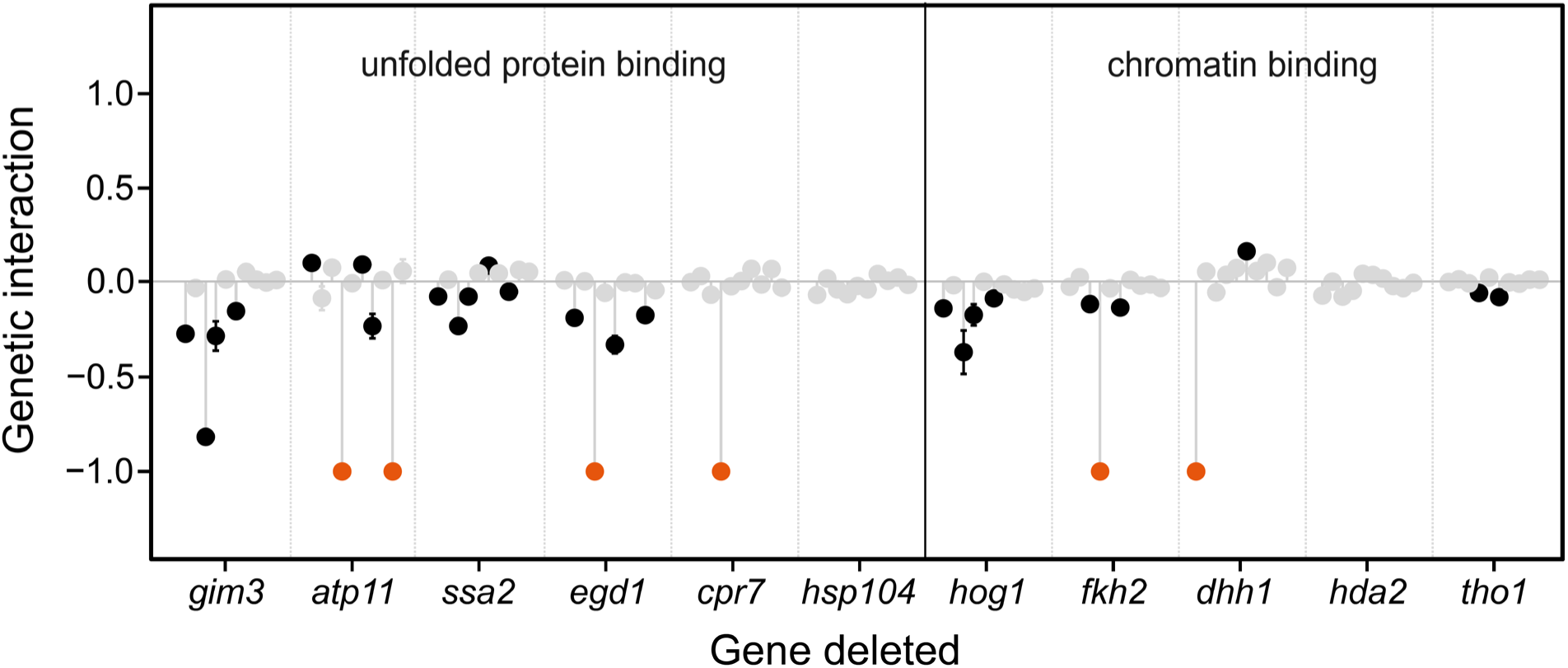
Epistasis between candidate buffer genes and random mutations. The deletion of genes with functions related to unfolded protein binding and involved in chromatin binding often show negative epistasis (negative genetic interaction) with UV-induced random mutations. Mutations often have a stronger negative fitness effect in these deletion backgrounds (negative epistasis), indicating that their effect can be buffered by the functional buffer gene. Orange dots indicate combinations of mutations and gene deletions that could not be generated, suggesting that these combinations are lethal. Each of the candidate buffer genes were deleted in ten mutant strains containing random mutations (dots represent the same 10 mutagenized strains for each buffer gene deletion, in the same order). A genetic interaction value of zero indicates that the fitness effect of the combination of random mutations and the deletion of a buffer gene is additive (no epistasis). Negative genetic interaction values indicate stronger negative effects than the additive effect (negative epistasis), positive interaction values indicate a less severe effect than the additive fitness effect of the random mutations and the respective gene deletion. Significant deviations from the additive model are in black (Wilcoxon rank sum test, *P*<0.01, error bars = se, *n* = 12 each), grey dots are not significant (Wilcoxon rank sum test, *P*>0.01, error bars = se, *n* = 12 each).

To obtain a more comprehensive view on the buffering potential, we continued the next experiment with five candidate buffer genes (*GIM3, SSA2, HSP104, HOG1, FKH2*). We measured, for each buffer gene, epistasis in a set of 200 randomly mutagenized strains (EMS mutagenesis; see Methods & Figure 4A). A large subset of mutagenized strains did show more severe fitness defects when the genetic buffer was removed (Figure 4B & Figure S3). Hence, the fitness of the double mutants was, on average, significantly lower compared to the expected fitness calculated by an additive model (Paired Wilcoxon signed rank test: *V* = 124750, *P* < 2.2 x 10^-16^). In addition, epistasis was tested for *HSP82*, and we found in agreement with our previous experiments, a smaller buffering potential (i.e. less negative epistasis; Figure S3). We further split up the data into three categories based on the initial fitness of the mutagenized strains (with the genetic buffer intact). Even without removing a genetic buffer, most of the introduced mutations (or set of mutations) were deleterious (Figure 4C; ‘Negative’), some mutations were neutral (Figure 4C; ‘Neutral’), and a small subset showed a slight positive effect (Figure 4C; ‘Positive’). Importantly, some mutations that were neutral in the presence of the buffer gene became deleterious when the gene was lost, indicating that at least part of the tested mutations were cryptic due to the activity of the candidate buffer genes.

**Figure 4.**
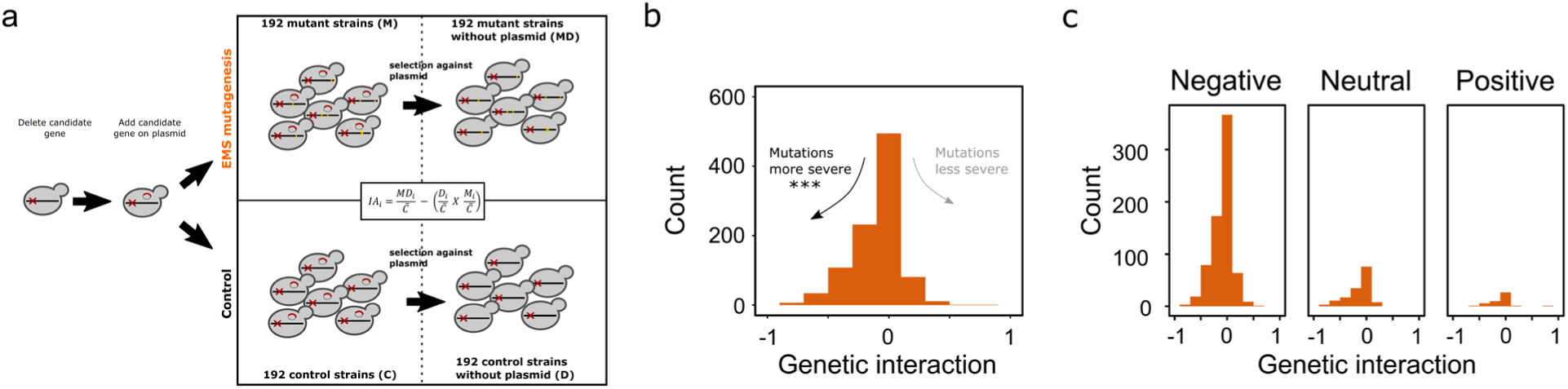
Large scale epistasis test. (A) Schematic representation of high throughput testing for epistasis between buffer genes and random mutations. For each tested buffer gene, the candidate buffer gene was deleted, and then re-introduced on a selectable and counter-selectable plasmid. A culture of the strain was randomly mutagenized using EMS, and 192 mutant strains were isolated. The fitness of mutant strains was measured when the plasmid was present (‘M’), and also when the plasmid was absent (after selection against the plasmid, ‘MD’). The same was done for a control culture, resulting in average fitness of 192 control strains with plasmid (‘C’) and average fitness of 192 control strains without the plasmid (‘D’), allowing us to calculate genetic interaction score (‘IA’) for each of the 192 mutant strains. This high throughput test was performed for five candidate buffer genes (*GIM3*, *SSA2*, *HSP104*, *HOG1*, *FKH2*), and the results of all tested buffer genes are represented together in panel b and c. (B) Deletion of a candidate buffer gene often results in more severe negative fitness effects of mutations (negative genetic interaction), while only a smaller subset shows less severe fitness effects of mutations (positive genetic interactions). Over all candidate buffer genes and mutant strains tested, mutations cause significantly stronger fitness effects when the buffer gene is removed (Paired Wilcoxon signed rank test, *V*=124750, *P*<2.2×10^-16^). (C) Data from panel b split based on the initial fitness of mutant strains (‘M’). The initial fitness of mutant strains (with the genetic buffer intact) is mostly negative, meaning that the mutations caused a decrease in fitness. A smaller subset is neutral (mutations have no effect on fitness) or positive (mutations cause an increase in fitness). Mutant strains were classified as neutral when mutant relative fitness was not significantly different from 1 (One sample t-test *, P*>0.05, *n* = 4 each), negative or positive when mutant fitness was significantly different from 1 (One sample t-test *, P*<0.05, *n* = 4 each). The deletion of a candidate buffer gene often results in more severe fitness effects of mutations (negative genetic interaction), regardless of the initial fitness of the mutant strains. Importantly, some mutations that have no effect on fitness (neutral), do show an effect when the buffer gene is absent, indicating that the effect of mutations could be buffered by the gene.

### Genetic buffers affect the maintenance and phenotypic expression of standing genetic variation

The findings in the present research showed that genes identified as genetic buffers showed extensive negative epistasis with *de novo* random mutations. However, the question remains to what extent such buffers also influence the phenotypic expression of standing genetic variation that has been subjected to purifying selection. To investigate this, we used a highly heterozygous feral yeast strain (Gallone et al., 2016) (South American isolate, ∼35000 heterozygous sites). Since this is a diploid yeast strain, different alleles can coexist in a heterozygote state, and recessive deleterious alleles can be retained. We sporulated this strain and subsequently dissected the tetrads. This was done for the WT, and two strains where we removed a genetic buffer by deleting a candidate gene belonging to either of the previously identified two GO categories (*HOG1* and *GIM3*). Interestingly, spore viability was lower in the latter two strains (viability: WT = 82%, *hog1* = 74%, *gim3* = 65%; Figure 5A), suggesting that more haploid genotypes were lethal when one of the genetic buffer genes was deleted. We then measured fitness of all the viable haploid spores and compared their fitness relative to the fitness of their corresponding diploid ancestral strain. For all three strains, the average fitness of the haploid segregants was lower compared to their diploid parent (Figure 5B, relative fitness <1). This is in agreement with previous observations of haploid segregants often showing reduced fitness compared to their diploid parent (Voordeckers et al., 2015). When comparing between the three strains, we did not find a significant difference in average fitness of the viable haploid segregants. Remarkably, when measuring the fitness of segregants with a buffer gene deleted in a stress environment (high temperature), a small number of haploid segregants showed a significant relative fitness increase (Figure 5B), which was not observed in the WT segregants. This does not only hint at the complex interaction between the buffering effect and the environment, but also suggests that deleting a buffer gene can in some cases expose beneficial cryptic variation.

**Figure 5.**
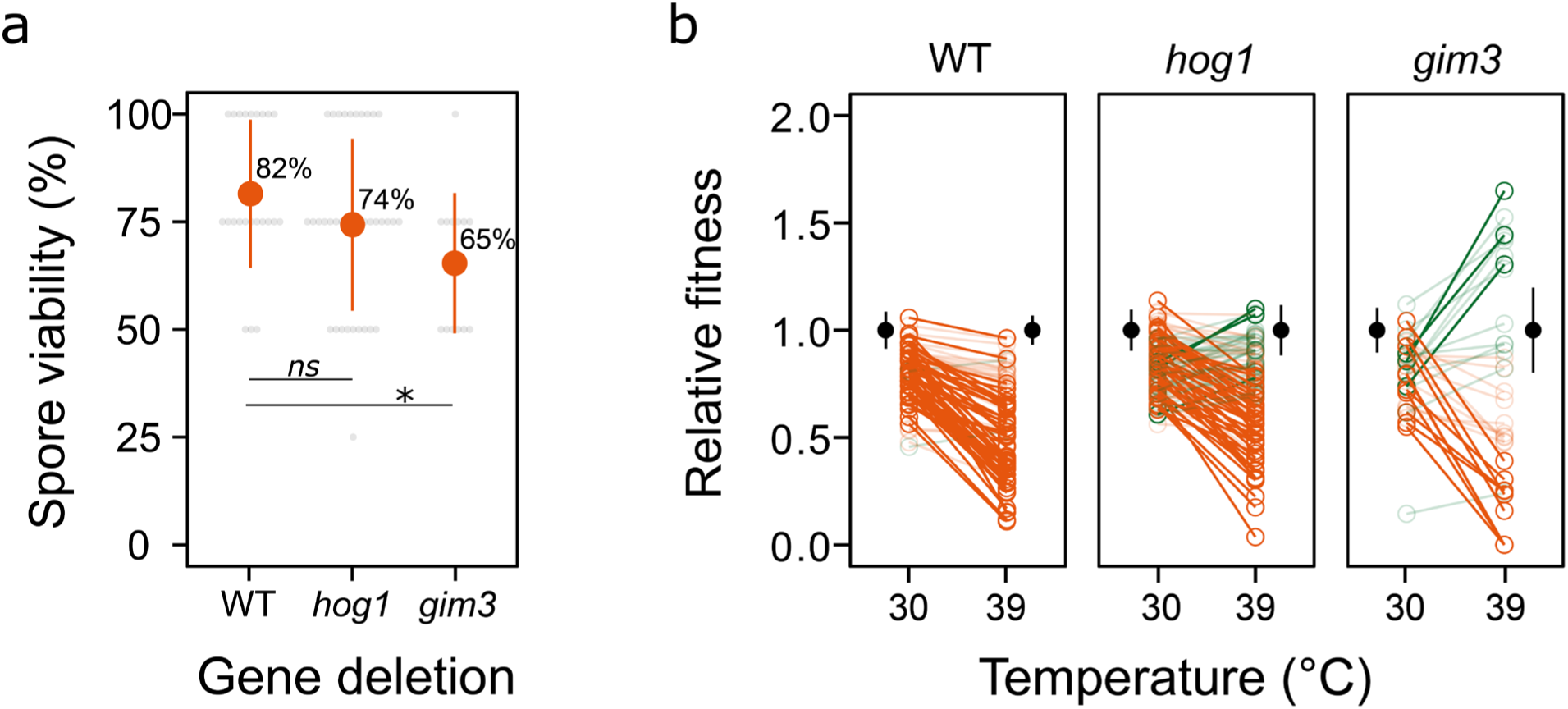
Genetic buffering potential in a natural yeast strain. (A) Spore viability after tetrad dissection of a WT strain (*n* = 23 tetrads dissected), and two strains with a candidate buffer gene deleted (*hog1*: *n* = 37 tetrads; *gim3*: *n* = 13 tetrads). Grey dots show the percentage viable segregants per dissected tetrad, orange dots show the average (+/- sd). Spore viability is lower when a buffer gene is deleted, but only *gim3* shows significant difference from WT (GLM with family = binomial, *gim3*: *z* = -1.285, *P* = 0.0324*, hog1*: z = -2.14, *P* = 0.1989). (B) Fitness of segregants (the viable spores from panel a) measured at different temperatures. Fitness is relative to the fitness of the diploid parental strain (measured at 30°C or 39°C, black dots +/- se) with no gene deleted (WT), or a candidate buffer gene deleted (*hog1* or *gim3*). Identical segregants are connected with a line (red = decrease in relative fitness, green = increase in relative fitness). Transparent lines when no significant difference in relative fitness between the two temperatures, opaque lines when significant difference in relative fitness (linear model, see methods, *n* = 3 each).

Based on these findings, we asked whether the presence of buffer genes affects the extent to which (cryptic) standing genetic variation is retained. The previously used diploid parent strains (WT, *gim3*, *hog1*) were subjected to two rounds of sporulation and random mating. After each sporulation round, we sequenced pools of randomly picked spores (500 spores per pool). This allowed calculating the frequency at which genetic heterogeneity (at previous heterozygous sites) was lost from the pooled spores (Figure 6A). On average, over all previously heterozygous sites, more heterogeneity was lost in strains lacking a genetic buffer (Figure 6B). The frequency of alleles at previous heterozygous sites changed stronger compared to the WT, with the effect being even stronger after the second round of sporulation (Figure 6C). Therefore, the number of alleles that were lost from the pools was significantly higher in both strains lacking a genetic buffer compared to the WT (GLM, family = quasipoisson, F*_(2,8)_* = 67.173, P = 7.814e-05; *gim3*, *t =* 7.708, *P =* 0.00025*; hog1*, *t =* 10.301, *P =* 4.89e-05). In other words, when a genetic buffer was removed, less (standing) variation was retained in the sequenced pools after meiotic cycles.

**Figure 6.**
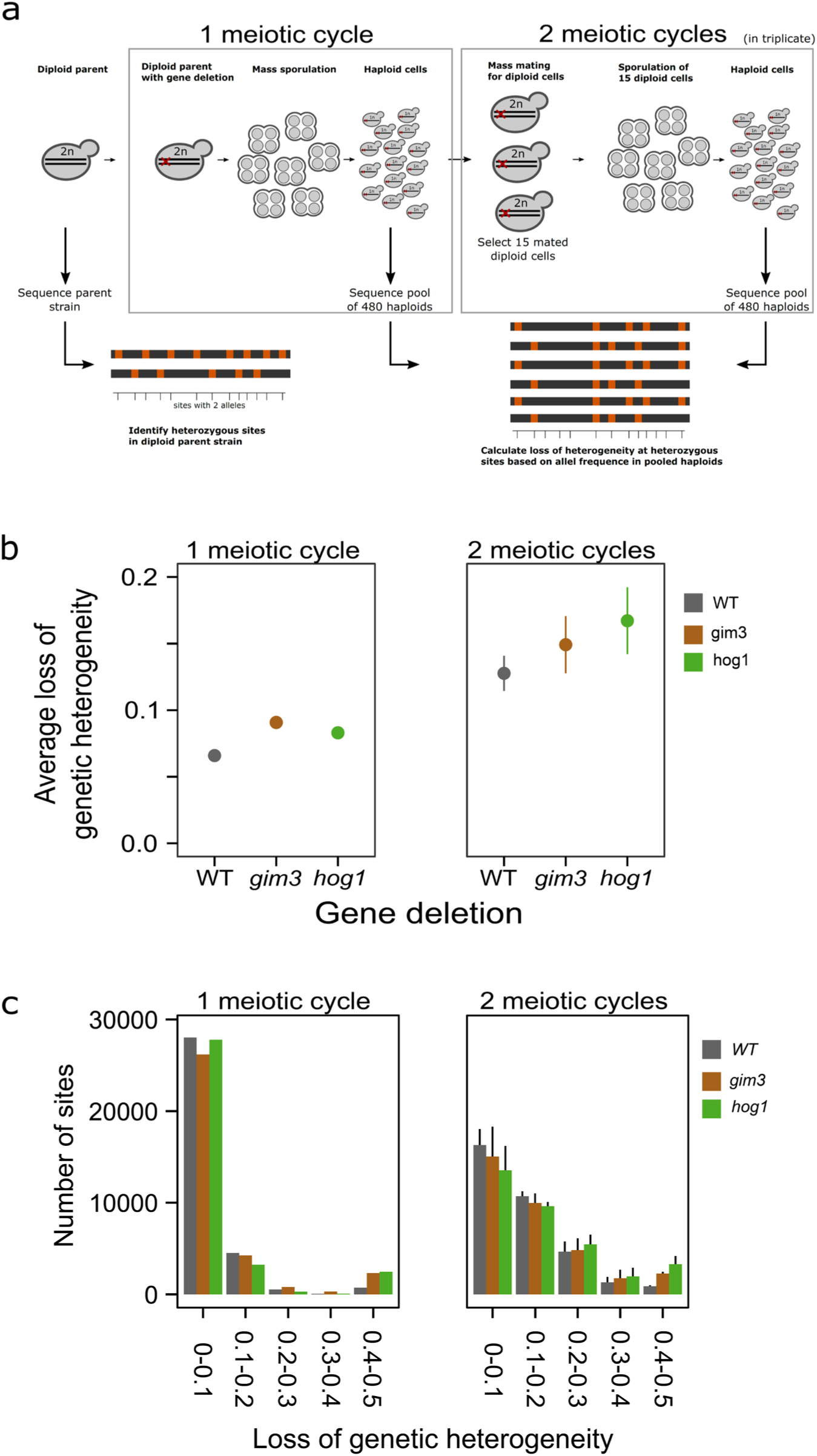
Loss of heterogeneity upon several meiotic cycles. (A) Schematic representation of meiotic rounds and sequencing to calculate loss of heterogeneity (or loss of standing genetic variation). From the diploid parent strain, a candidate gene (*GIM3* or *HOG1*) was deleted. This train was sporulated, resulting in millions of tetrads, and subsequently even more haploid cells (after removing spores from tetrads). The pool of haploid cells was used for random mating, forming diploid cells again. 15 diploid cells were selected for usage in another round of sporulation, resulting in a second pool of haploid cells (selection of diploids and second round of sporulation was performed in triplicate). The Diploid parent stain was sequenced to identify heterozygous sites (color represents different alleles). From the first pool of haploids (first meiotic round), 480 haploids were isolated and sequenced as a pool. Also 480 haploids were isolated from the second meiotic round, and sequenced as a pool (three pools, since there were triplicates). From all pools, allele frequency at previously identified heterozygous sites was calculated, allowing us to investigate the loss or fixation of alleles, and the change in allele frequency at these previously heterozygous sites. (B) Average loss of genetic heterogeneity for each strain after one (1 replicate) or two meiotic cycles (3 replicates, +/- sd). Loss of genetic heterogeneity was calculated by taking the average change in allele frequency from the pooled spores at previous heterozygous sites (identified in the diploid parental strain). (C) The change in allele frequency at previously heterozygous sites, calculated from the pooled spores upon one meiotic cycle (cycle 1, one replicate) and two meiotic cycles (cycle 2, three replicates; +/- sd). More alleles are (nearly) completely lost (change in allele frequency = 0.4-0.5) when a candidate buffer gene is deleted (*gim3*, *hog1*).

## DISCUSSION

We used the model eukaryote *Saccharomyces cerevisiae* to identify genes that act as genetic buffers, i.e. genes that influence the fitness effect of a relatively large fraction of mutations. We found two GO categories, ‘unfolded protein binding’ and ‘chromatin binding’ that showed an enrichment for genes with a strong buffering potential (Figure 2A). The first category contains chaperone genes like *HSP90*, the seminal previously studied example of a genetic buffer gene (Zabinsky et al., 2019). Note that in our screen, *HSP90* was not a top candidate likely because of redundancy by its paralog. The second category of genes, involved in ‘chromatin binding’, is also of particular interest. This category contains genes involved in cellular regulation, including several chromatin regulators. We suspect that these genes can compensate mutations by altering gene expression (Lehner et al., 2006; Tirosh et al., 2010). Genes within the two identified GO categories, and especially the top candidate genes, interact with many other genes, indicating that they are central in the genetic interaction map (hub genes) and can act as modifiers of genetic variation. This agrees nicely with the definition of a buffer gene – reducing the activity of a buffer changes the fitness effect of many random mutations. It is worth noting that genetic interaction screens differ from our screen for buffer genes. Previous screens to map genetic interactions typically measured epistasis between two genes (by deleting both genes), instead of between a gene and random (point) mutations. In this way, these screens did not quantify the crucial interaction between mutations and potential buffering functions of genes.

The findings in this study provide insights in the characteristics and functions of genetic buffers. They tend to be relatively highly connected within the genetic network and often function in protein folding (chaperone proteins) or are involved in chromatin regulation. Previous work regarding *HSP90* suggested a straightforward buffering mechanism; chaperone proteins could help mutated proteins to fold correctly, thereby maintaining their function (Zabinsky et al., 2019). Many of our top candidate buffer genes are directly involved in protein folding. Furthermore, the regulation of compensatory pathways through chromatin structure points toward a functional explanation for chromatin regulators as buffers, by helping to mitigate perturbed gene expression patterns. However, these mechanistic explanations are not necessarily the only way to provide genetic buffering, most importantly when considering fitness effects of mutations. For example, negative fitness effects of unfolded proteins could be decreased by clearing potentially harmful misfolded proteins, thereby preventing proteotoxicity, rather than folding misfolded proteins directly. Interestingly, the GO category ‘exocytosis’ also appears to display significant buffering function (Figure 2A).

One of our strongest candidate genes (*GIM3*) is a highly connected hub gene (Lehner et al., 2006; Tong et al., 2004) and is a subunit of the prefoldin complex, which assists folding of cytoskeleton proteins. Hence, *GIM3* may buffer genetic variation by directly interacting with these proteins, but also more generally, by its involvement in the maintenance of protein homeostasis or stress induced transcription (Tahmaz et al., 2022; Valiadi et al., 2012). *HOG1* is another strong candidate gene in our study, which is a hub gene belonging to the chromatin binding GO category. *HOG1* is a mitogen activated protein kinase, that mediates gene expression of 600 genes, often through interactions with chromatin modifiers, allowing cells to cope with stressful conditions (Tirosh et al., 2010; Westfall et al., 2004). Therefore, *HOG1* could function as genetic buffer, by maintaining robust biological outcomes despite mutations, or more indirectly by its general role in stress response. Interestingly, physical and genetic interactions with *HSP90* have been reported for some candidate genes such as *GIM3* and *HOG1* (Girstmair et al., 2019; Millson et al., 2005; Yang et al., 2006; Zhao et al., 2005), indicating that they could be client proteins of *HSP90*, and therefore be part of a buffering network. Thus, the exact underlying mechanisms may vary for each specific case and could be the result of a combination of different functions. The elucidation of the exact mechanism by which genetic buffers act would require further in-depth characterization through, for instance, specific phenotyping assays.

Our experiments have shown that genetic buffers can affect the fitness effect of a broad range of *de novo* mutations. However, buffer genes do not influence the effect of all mutations, and different genes may interact with different subsets of mutations. The fitness effect of many *de novo* mutations with the genetic buffer intact was deleterious (Figure 4C), indicating that genetic buffers often do not fully suppress the deleterious fitness effect of *de novo* mutations. Importantly, however, a small fraction of mutations were neutral when a buffer gene was present, but became non-neutral upon deletion of a specific buffer gene (Figure 4C), which is in line with the classic definition of a genetic buffer. This observation indicates a potential role of genetic buffers in CGV, but also supports previous hypothesis that truly cryptic mutations might be rare, and therefore only slowly accumulate in the genome over time (Geiler-Samerotte et al., 2016; Richardson et al., 2013; Siegal and Leu, 2014). However, given that many genes could possess a buffering function (Figure 2), the potential of CGV regulated by the entire genetic network may be substantial. In a natural yeast strain, we found that more alleles proved incompatible with a successful meiotic cycle when the genetic buffer was removed (Figure 5A), and more genetic variation was maintained in the presence of genetic buffers (Figure 6B & C). Hence, some of the natural standing variation depends on the presence of specific genetic buffers, and genetic buffers may thus be vital for the accumulation and maintenance of standing genetic variation, including CGV.

In this study, we mostly focused on the buffering effect based on fitness rather than specific phenotypes. The question of how our candidate buffer genes influence the variation in more specific phenotypes demands further investigation. The effects observed when cells were subjected to temperature stress hints that buffer genes affect phenotypes like stress resistance. We also found evidence for natural cryptic variation that depended on the interaction of the genetic buffer and the environment (Fig 5b), illustrating that in a new environment, some cryptic variation could provide an adaptive benefit. However, this depended on the removal of the genetic buffer, which often comes at a fitness cost by itself. The evolutionary implications need to be further investigated and more directed experiments are needed to investigate how important this phenomenon is in more natural conditions. All our top candidate genes are relatively conserved genes and have orthologs in many different organisms (from human to drosophila), strengthening the idea that buffer genes are important for complex genetic diseases such as cancer (Siegal, 2017). Furthermore, some of our identified buffer genes have been linked to neurodegenerative diseases (e.g. *SSA2*, prefoldin, *GIM3*: Broer et al., 2011; Liang et al., 2020, and *Hsp104*: March et al., 2020), suggesting a possible link between modifiers of genetic variation and disease state.

## ACKNOWLEDGMENTS

We thank all K.J.V. laboratory members for their suggestions. We also thank Valerie Neven for her help and suggestions. We specifically thank Peter Bircham for his suggestions regarding fitness experiments. This work was supported by a postdoctoral fellowship from Research Foundation Flanders (FWO) to J.F. (12P7219N), an FWO PhD fellowship to M.T.T. (11H1823N) and S.B. (11PS824N), an FWO junior research project (G012524N) and an FWO ERC runner-up project (G0E1223N) to S.C.V. and a KU Leuven C1 grant (C16/23/007) to K.J.V.

## AUTHOR CONTRIBUTIONS

J.F., S.C.V. and K.J.V. conceived and designed the study and experiments with input from D.F.J.. J.F., M.T.T., E.B., S.B., J.R., K.V. performed the experiments and collected and analyzed the data. J.F., M.T.T, K.V. and K.J.V. wrote the manuscript and all authors contributed to the final version of the manuscript.

## METHODS

Methods will be formatted to STAR methods.

### Data and code availability

Raw sequence reads from genomic analyses are deposited in the Sequence Read Archive with accession number PRJNA837038. Other data reported in the paper are available in Supplementary information as mentioned in Methods. Any additional information required to reanalyze the data reported in this paper is available from the lead contact upon request.

### Genome-wide screen

#### Setup

Using the haploid yeast single-gene knockout collection (YKO: Giaever et al., 2002; Winzeler et al., 1999, *MAT* **a**), we measured the fitness effect of random mutations in each of the deletion strains. Each strain of the YKO was cultured overnight in 96 well plates in 200 μl SCM (synthetic complete medium; Table S3) by pinning (using singer rotor HDA) from -80°C stock cultures (96 well format containing 20% glycerol). Overnight cultures were diluted (800x) and split into a ‘control culture’ and a ‘mutant culture’ (40 μl each). The mutant culture was exposed to UV-light (9000 x 100 uJ/cm^2^, CL1000 ultraviolet crosslinker UVP) to introduce different random mutations in each cell. Mutations were fixed by growing the cultures (40 μl culture + 125 μl SCM, for control and mutant culture) in the dark (20°C, ∼2 generations). Cultures were then diluted in SCM (mutant culture: 12x, control culture 36x) and seeded into 384 well plates (384-well glass bottom microwell plate, Brooks) coated with ConA (Concanavalin A, Canavalia ensiformis, Sigma Aldrich). Using automated microscopy (inverted Nikon Eclipse Ti microscope with a DL-604M-#VP camera; AndorTM technology, 20X/0.75 Nikon lens), micrographs were taken every hour from 5 positions within each well (software: Metamorph 7.8.0.0; Molecular Device, LLC; 12h, 30°C). The micrographs were processed with Cellprofiler 2 (Carpenter et al., 2006, version: 2.2.0) in order to identify and track micro-colony size over time. These were converted into growth curves for each of the tracked cells in R (R Core Team, 2021) with Rstudio (Rstudio, 2021) and fitness of each tracked cell was calculated by taking the slope of the log-transformed growth curves. Micro-colonies were only processed when identified before frame 4, and when tracked for minimum 3 successive frames. We only included deletion strains that had at least 50 individual cells tracked in the control and in the mutant culture. To generate a measure of buffering potential, we compare the (average) fitness of the cells in each mutant culture relative to the (average) fitness of cells in the corresponding control culture (Table S4). Finally, we removed deletion strains from which the deleted gene functions in DNA repair (resulting in a final dataset of 3484 deletion strains). Such strains would accumulate more mutations, and therefore have stronger fitness defects due to more mutations and not due to a potential buffering effect.

#### Processing results

Pseudogenes (dubious ORFs) were identified using the genome annotation file (https://downloads.yeastgenome.org/sequence/S288C_reference/genome_releases/S288C_refere nce_genome_Current_Release.tgz [accessed: 2020-03-23]). The genome annotation file was also used to calculate the percentage overlap between overlapping ORFs, and we only used overlapping ORFs when overlap was >20%.

For each deletion strain used in our analysis, we search for all their corresponding GO terms (of the deleted gene, excluding gene ontologies belonging to cellular component group). Gene associations were downloaded froms SGD (http://sgd-archive.yeastgenome.org/curation/literature/gene_association.sgd.gaf.gz [GO version: 2017-08-27]). Genes related to DNA repair were removed based on all GO terms in the database containing ‘repair’ and ‘damage’ (resulting in 3484 deletion strains in total). We restricted our analysis to GO terms that had at least 20 genes associated with them (thus, most deletion strains appear in multiple GO categories, because they are associated with multiple GO terms). The average relative fitness of all genes for each GO term was calculated and is represented in figure 2A and Table S5.

We performed a GO enrichment analysis on our dataset, and searched for enriched GO terms by ranking genes according to their buffering potential (Wilcoxon rank-sum test for enrichment). We restricted the GO enrichment analysis by only including GO terms that had at least 20 genes associated with them, to comply with figure 2A. GO enrichment analysis was performed using the GOfuncR package (Grote, 2021, version 1.14.0) and results of the enrichment were corrected for multiple testing (Table S2).

We tested whether there was a relationship between our measure of buffering potential and known genetic interaction properties of the tested genes, which are archived in the BioGrid database (Stark et al., 2006, *Saccharomyces cerevisiae*_S288c-3.5.174). The BioGRID dataset is a public database collecting genetic and protein interaction data from model organisms. Relative average fitness was used as a dependent variable, and the different interaction types as independent variables in a Generalized Linear Model (GLM, family = Gamma, link = log). Stepwise model selection was used to select the model with significant predictors and influential observations were removed based on cooks distance. Out of the 11 different genetic interaction types archived in the BioGRID database, two showed a significant correlation with our measure of buffering potential (Table S1).

We counted the number of genetic interactions for each gene by using the SGA NxN dataset (Costanzo et al., 2016). For each gene, the number of significant interactors was counted based on p-value < 0.05. We tested for significant more interactors between genes contained in the ‘unfolded protein binding’ and ‘chromatin binding’ GO category, and all other genes (deletion strains) in YKO using GLM (GLM, family = quasipoisson, link = log).

### Identification of mutations introduced by UV exposure

To test if UV exposure resulted in different random mutations in each cell, we sequenced the genomes of 10 isolates from a mutant culture and 10 isolates from a control culture (using the YKO background strain). Cultures were plated on SCM agar (SCM + 2% agar) and 10 colonies were randomly picked per culture, and one isolate of the S288C background strain was included (‘ancestor’). DNA was extracted from each isolate (QIAGEN Genomic-tip 20/g) and sequenced using Illumina Hiseq 150 bp paired end sequencing (raw sequencing reads are deposited at Sequence Read Archive; PRJNA837038). The raw reads were trimmed using Trimmomatic (Bolger et al., 2014) (version 0.38) and mapped to the reference genome (S288C_reference_sequence_R64-2-1_20150113.fasta, http://sgd-archive.yeastgenome.org/) using bwa mem (Li and Durbin, 2009, version 0.7.12-r1039). Picard (version 2.18.12, http://broadinstitute.github.io/picard/) was used to mark duplicates and to add read group information. Variant calling was done with GATK (McKenna et al., 2010) (version 3.8-1-0-gf15c1c3ef) following their standard practice guidelines (haplotypecaller and genotypeGVCFs). The variant file was converted to table format for further processing in R. Single nucleotide variants were quality filtered (DP>100, FS<40, MQ>40, QUAL>40, QD>5) and INDELs removed. Variants identified in the ancestor were removed as they already existed before UV treatment. SnpEff (Cingolani et al., 2012) was used to annotate the variants to determine their impact. For the UV treatment and control, we also sequenced pools of 100 isolates (two pools each). Individual colonies (100 per pool) were picked from the SCM agar plates (plated cultures above) and grown overnight in 96 well plates. Cell densities were adjusted to equal densities based on OD, and the 100 isolates for each pool were mixed together for DNA isolation (QIAGEN Genomic-tip 100/g). Sequencing was done as described above, with ploidy set to 100 and variant were quality filtered (DP>2000, FS<40, MQ>40, MQRankSum>-10, ReadPosRankSum>-8, QUAL>50), keeping only variants with frequency > 0.005.

### Candidate genes and strains

We selected 11 candidate genes for further testing (Figure 2B in bold). We only selected candidate genes that were annotated for the ‘unfolded protein binding’ or ‘chromatin binding’ function manually, in order to have a conservative selection of candidate genes.

#### Construction candidate gene strains

All (deletion) strains used for further testing and validations in further experiments were constructed from the same parental WT strain (BY4741, *MAT* **a** his3Δ1 leu2Δ0 met15Δ0 ura3Δ0) and candidate genes were deleted using the standard yeast LiOAc transformation protocol and homologues recombination, where the candidate gene (ORF) was replaced with the Hygromycin marker (Table S3).

#### Fluctuation assays

To determine mutation rates, we performed fluctuation assays following Voordeckers et al., 2020. For each candidate deletion strain, a single colony was used to inoculate a culture in 5ml SCM (30°C). After one overnight growth, each culture was diluted 5000x and used to inoculate 4 rows of a 96 well plate. After one overnight growth, all wells from one row were pooled and used to measure cell density using cellcounter (TC20 Automated cell counter, Biorad). Each well of the next 3 rows was plated on SCM-Arg with canavanine (60 mg/L canavanine, Sigma Aldrich, Table S3), and colonies were counted after 3 days of incubation (30°C). The cell densities and the number of colonies on canavanine plates were used to calculate the mutation rate using the R package rSalvador (Zheng, 2017) using Maximum likelihood estimation of the mutation rate accounting for variation in Nt; Table S1.

#### UV sensitivity assays

The candidate deletion strains were inoculated in 96 well plates in SCM and serial diluted 8 times (2x dilution each). The serial dilution allowed us to take exponential growing cultures after overnight incubation (30°C) by selecting OD<1.0 (Tecan infinite M200 Pro). Each culture was then diluted to 1.5 x 10^5^ cells/mL, which was used for a 2x dilution series (8 dilutions in total). 10 μl of each dilution was spotted on YPD agar plates (2% agar; Table S3) and exposed to 0, 25 or 50 uJ/cm^2^ of UV light. Plates were scanned after 3 days of incubation (in dark, 30°C). For this assay, two positive control strains were included where a gene involved in UV sensitivity was deleted (*RAD10* and *RAD52*; taken from YKO). Based on the spotting assay, the two positive control strains show clear UV sensitivity whereas the other candidate genes did not show a difference from the WT strain (Figure S2).

#### Epistasis assays

Ten randomly mutated strains were used in this assay. These strains were the same as the 10 mutant strains used for sequencing (see above section: ‘Identification of mutations introduced by UV exposure’). Each of the candidate genes were deleted separately in each of these mutant strains (resulting in 110 strains). We measured the fitness of each of these strains (‘MD’; random Mutations and gene Deletion), and also the fitness of strains where the candidate genes were deleted in the WT background strain (11 strains; ‘D’; Deletion) and the fitness of each of the ten mutant strains (‘M’; random Mutations). Fitness was measured by spotting assays using the singer rotor HDA as follows: all strains were stored in -80°C (in SCM + 20% glycerol) in 96 well plates. Frozen plates were thawed and used to inoculate new 96 well plates containing SCM (by pinning with singer rotor HDA). After one overnight (30°C), 96 well cultures were combined together by pinning on SCM agar (density of 384, edges and empty spots were filled with BY4741 to eliminate edge effects). After 48h incubation (30°C), the agar plate was re-arrayed to a density of 1536, resulting in four technical replicate spots from each source spot and incubated for 48h (30°C). We repeated this procedure twelve times to generate twelve independent replicates. All plates were scanned and we used the gitter R package (Wagih and Parts, 2014) to measure the area of each spot (colony) as a proxy for fitness. All fitness measurements were normalized to a wild-type BY4741 included in each plate to obtain relative fitness values. Next, we calculated genetic interaction score (epistasis) following Wagih et al., 2013. This score reflects whether the measured fitness of MD-strains (strains containing a gene deletion and random mutations), deviates from the predicted effect, calculated from the fitness of D-strains (strains only containing a gene deletion) and M strains (strains only containing random mutations). Thus, genetic interaction = MD – (M x D). We tested for significant epistasis by testing whether M x D was significantly different from MD (Wilcoxon rank sum test, Table S6).

For the large scale epistasis test, candidate buffer genes were deleted from the genome (cf. ‘*Construction candidate strains’*, BY4741 background), resulting in 6 strains, each having one gene deleted (*GIM3*, *SSA2*, *HSP104*, *HOG1*, *FKH2*, *HSP82*). Next, for each of these strains, the deleted candidate buffer gene (+200 bp upstream and downstream) was transformed on a plasmid containing the URA marker and transformed back in the deletion strain (Table S3). Thus, the 6 resulting strains each have one gene moved from the genome to a selectable and counter-selectable plasmid. The strains were grown overnight in 5 mL SCM-Ura (Table S3), and 1 ml of the cultures was washed with sterile water and resuspended in 1ml 0.1M sodium phosphate buffer containing 30 μl EMS (Ethyl methanesulfonate, Sigma Aldrich). After 1h of incubation (30°C), the cells were washed with 5% sodium thiosulfate buffer and resuspended in 1 ml SCM-Ura. The resulting mutant cultures were plated for single colony selection (SCM-Ura + 2% agar). For each strain, a control culture was established in the same way, but EMS was replaced by water. 192 mutant colonies and 192 control colonies were picked, and grown for two overnights (30°C) in 96 well plates containing 150 μl YPD. These cultures were pinned on agar plates using singer rotor HDA to a density of 384 on (I) SCM-Ura agar (selection for the plasmid), and (II) SCM +5FOA agar (selection against the plasmid; Table S3, 2% agar). This counter-selection resulted in a culture without the plasmid, and therefore loss of expression of the candidate buffer gene. We verified loss of gene expression by qPCR for four strains (WT, *GIM3*, *HOG1*, *HSP82*; Figure S4) using Lucigen Masterpure Yeast RNA Purification Kit for RNA extraction and reverse transcription, followed by qPCR (AB, step on plus) using SYBR green (PowerSYBR green PCR Master Mix, Applied Biosystems), *TDH3* as endogenous control and the WT strain for relative quantification (Table S3). The agar plates from (I) and (II) were incubated (30°C) for 48h and re-arrayed to a density of 1536 on SCM-Ura or SCM+5FOA respectively, resulting in four replicate spots from each source spot after incubation (48h, 30°C). The resulting colonies were pinned on SCM agar plates, and plates were scanned after 48h incubation (30°C). Genetic interaction scores were calculated as described above, where MD = mutant strains from SCM+5FOA, M = mutant strains from SCM-Ura and D = control strain from SCM+5FOA (Table S7). For this assay, fitness values were calculated by taking colony sizes relative to the average colony size of control strains on SCM-Ura. We tested whether mutant strains had a significant different fitness by testing whether the relative fitness of mutant strains (M, four replicates) was significantly different from 1 (one sample t-test, Table S7). Before each t-test, we tested for normality. Samples that did not meet the normality assumption were excluded from the analysis in Figure 4B (∼5% of samples).

### Spore dissection

Three strains were used for spore dissection. First, the HO locus from yeast strain BI002 (Gallone et al., 2016) was deleted (= ‘WT’ strain, Crispr-cas9 transformation following Vyas et al., 2018 and Table S3), and two additional strains were made by additionally deleting a candidate buffer gene in the HO deletion strain (*GIM3* or *HOG1* respectively, Cripsr-cas9 transformation; Table S3). Each strain was grown overnight in 5 ml YPD, from which 1 ml was used to inoculate 50 ml of pre-sporulation medium (8 g/l yeast extract, 3 g/l peptone, 10% glucose). After 5h incubation (30°C), cells were washed with water, and OD adjusted to OD15. 20 μl was used to spot on MSM plates (Table S3), which were incubated for 7-10 days (23°C). From MSM plates, cells were treated with zymolase (Zymolase 100T, Amsbio) and resulting tetrads were dissected with dissection microscope (Singer instruments, MSM 400). Spores were validated by mating type PCR (Supplementary Table3). From each dissected tetrad (WT = 23, *gim3* = 13, *hog1* = 37 tetrads dissected), spore viability was determined by growth on YPD agar, and viable spores were stored in -80°C (96 well plates with YPD + 2% glycerol). The fitness of all viable spores was measured by pinning assays using the singer rotor HDA. 96 well plates were thawed from -80°C and used to inoculate new 96 well plates containing 120 μl YPD. After one overnight (30°C), 96 well cultures were pinned on SCM agar to a density of 384. After one overnight (30°C), colonies were pinned to new SCM agar plate, to a density of 1536 (4 technical replicates per spore) and incubated (14h). In this last step, colonies were pinned on one extra agar plate and incubated (14h) at 39°C. We repeated this procedure three times to generate three independent replicates. All plates were scanned and we used the gitter R package (Wagih and Parts, 2014) to measure the area of each spot (colony) as a proxy for fitness. All fitness measurements (colony size) were normalized to their diploid ancestor strain incubated at the same temperature on the same plate and for each, the average of 4 technical replicates was taken (e.g. colony of a spore with *GIM3* deletion, was normalized to the diploid strain with *GIM3* deletion; Table S8). We tested each spore for significant differences in fitness growing at 30 and 39 degree using the three biological replicates (LM after testing for normality Table S8 & Figure 5B, when non normal data, results were considered not significant).

### Loss of heterozygosity

The three strains generated above (‘Spore dissection’, diploid BI002 background; WT, *hog1*, *gim3*) were treated and spotted on MSM plates as described above (‘Spore dissection’, 40 μl per spot, 35 spots per plate and strain). After 12 incubation days (23°C), spots were washed from MSM plates with 10 ml water. 800 μl Zymolase (1 mg/ml) and 16 μl 2-Mercaptoethanol (Sigma Aldrich) was added and incubated overnight (35°C, 80 rpm). The cell suspension was transferred to 15 ml falcon tube containing sterile glass beads (425-600 um) and vortexed vigorously for 3 minutes. Suspension was washed with water and dissolved in 9 ml 0.75% Triton (Triton X-100, Sigma Aldrich), put on ice for 15 minutes and sonicated (3 x 30 seconds, 20% Amplitude). The resulting spores were washed and resuspended in 1 ml water. Mating type PCR was used to evaluated sporulation efficiency (98% haploid cells, n=192) by plating out spores on YPD and picking individual colonies. 50 ml GNA (Table S3) was inoculated with 5×10^5^ cells/ml washed spores and allowed for mating by overnight incubation at 30°C (80rpm). Cells were plated out on YPD and per strain, 3×15 diploid (mated) cells (confirmed by mating type PCR) were isolated. The 15 diploid cells were grown overnight in 200 ul, pooled and used as starting culture for the second round of sporulation. This second sporulation round was done using 3 replicates. For each strain and replicate, the washed spores from the first sporulation round and second sporulation round (in triplicate) were plated out on YPD, and of each, 480 individual colonies were picked, grown overnight in 200 μl YPD in 96 well plates.

For each strain and replicate, all 480 cultures were pooled, DNA was extracted (QIAGEN Genomic-tip 100/g) and sequenced (150 bp paired end sequencing, DNBseq; raw sequencing reads are deposited at Sequence Read Archive; PRJNA837038). Thus, we sequenced for each strain 1x pool of 480 isolates (first sporulation round; >1000x coverage), 3x pool of 480 isolates (second sporulation round, >1000x coverage). Additionally, we sequenced the diploid parental strain (BI002 with HO locus deleted; ∼200x coverage) in order to identify heterozygous sites. Variant calling was done as explained above, but GATK4 (McKenna et al., 2010) was used (version 4.2.2.0), and only SNPs were processed and quality filtered (QD>2, QUAL>30, SOR<3, FS<60, MQ>45, MQRankSum>-12.5, ReadPosRankSum >-8, GQ>20). Variant sites and number of mapped reads per allele were converted to table format and further processed in R. For each of the heterozygous sites identified in the diploid parental strain, we calculated the allele frequency of those alleles in the pooled samples, based on the number of mapped reads per allele. This allowed us to calculate the difference from pure heterozygosity (pure heterozygous = 0, one allele fixed = 0.5; Figure 6B). Furthermore, we counted the number of sites where one of the two alleles at previously heterozygous sites was completely lost from the pooled samples. This data was used to test for significant differences in lost alleles between the three strains at the second meiotic round.

### Statistics

Statistical analyses were performed in R (R Core Team, 2021) and Rstudio (Rstudio, 2021). Linear models and one sample t tests were used on normally distributed data. Two-sided Wilcoxon rank sum tests and GLM with appropriate family and link function were performed for non-parametric data. *P* values of 0.05 were used as the threshold for significance unless noted otherwise. Data is shown as mean +- standard deviation unless noted otherwise. The R package rSalvador (Zheng, 2017) using Maximum likelihood estimation of the mutation rate accounting for variation in Nt, was used to test for significance, *P* values were adjusted for multiple testing using Benjamini & Hochberg. GO enrichment was performed using the GOFuncR (Grote, 2021) package and to correct for multiple testing and interdependency of the tests, family-wise error rates were computed based on random permutations of the gene-associated variables.

## SUPPLEMENTARY INFORMATION

**Supplementary information** includes four figures and eight tables and can be found with this article online at DOI X.

**Supplementary Table 1**

Statistics for (I) correlation with BioGRID database and (II) Fluctuation assay.

**Supplementary Table 2**

Results from GO enrichment.

**Supplementary Table 3**

Overview of media used in the study, generation of deletion strains and plasmids, primers used for mating type pcr and qPCR.

**Supplementary Table 4**

Growth rates measured in the genome-wide screen.

**Supplementary Table 5**

Average growth rates of deletion strains grouped by GO category.

**Supplementary Table 6**

Fitness values, genetic interaction scores and *P* values for mutant strains and candidate gene deletions.

**Supplementary Table 7**

Data obtained in the high throughput epistasis assay. Fitness values, genetic interaction scores and *P* values (testing if mutant strains have a relative fitness different from 1) for mutant strains and candidate gene deletions.

**Supplementary Table 8**

Fitness values of spores measured at 30°C and 39°C, including *P* value for normality testing (Shapiro wilk test) and P value testing for difference in relative fitness between 30°C and 39 °C.

## Supplementary figures

**Supplementary figure 1.**
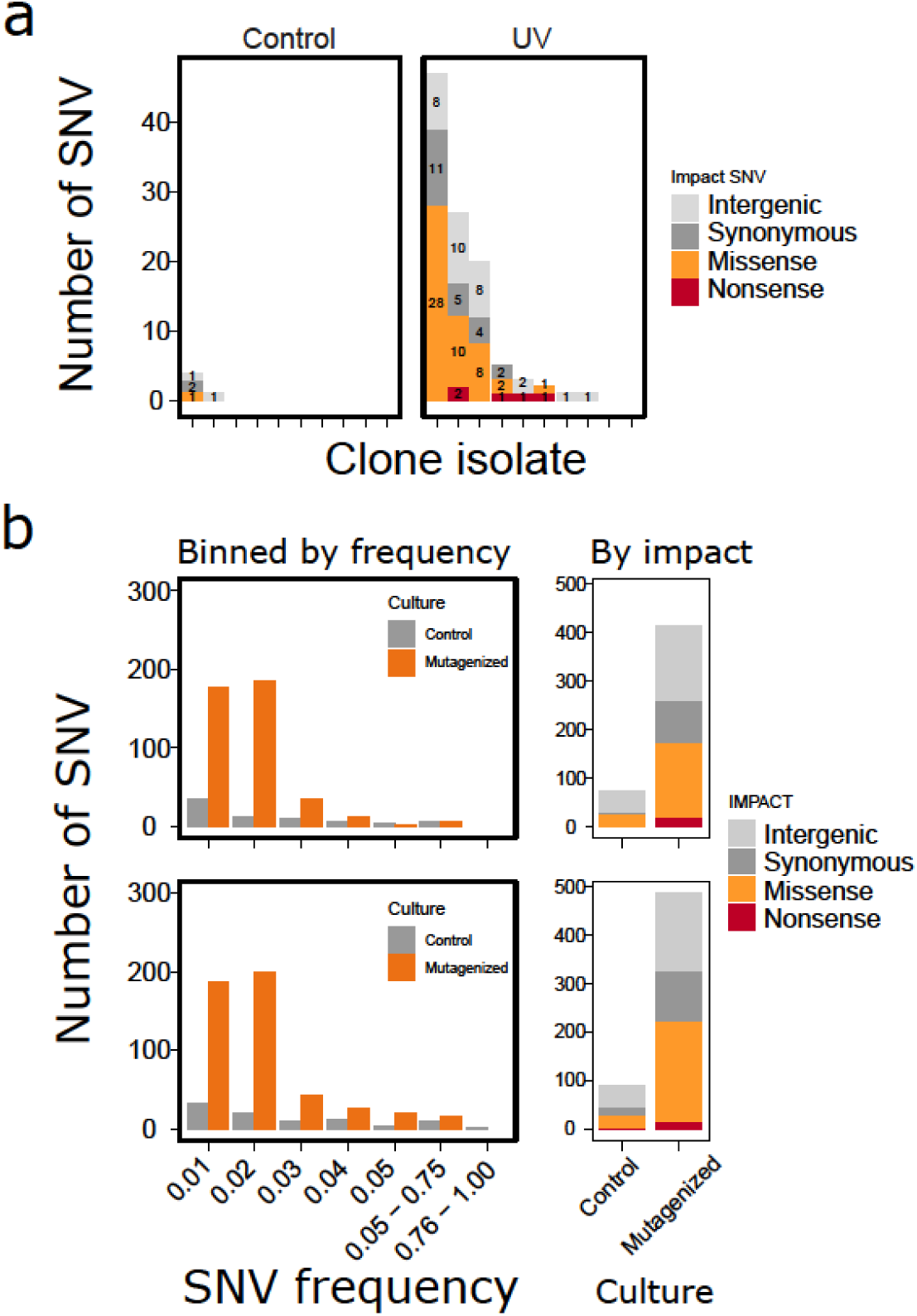
(A) Number of SNVs (single nucleotide variants) identified in 10 isolates from a control culture and 10 isolated from a mutagenized culture (UV mutagenesis). SNVs were divided based on the predicted effect of the mutation (intergenic variants, synonymous variants, missense variants or nonsense variants). (B) SNVs were identified in two pools of 100 isolates form two control cultures and two pools of 100 isolates from mutagenized cultures. SNVs were binned by the frequency in the pools, showing that most SNVs were at low frequency. Thus, the mutagenized cultures consisted out of cells with different mutations. Control pools showed less mutations compared to mutagenized cultures. The total number of SNVs was counted and divided based on the predicted effect of the mutations (cf. A).

**Supplementary figure 2.**
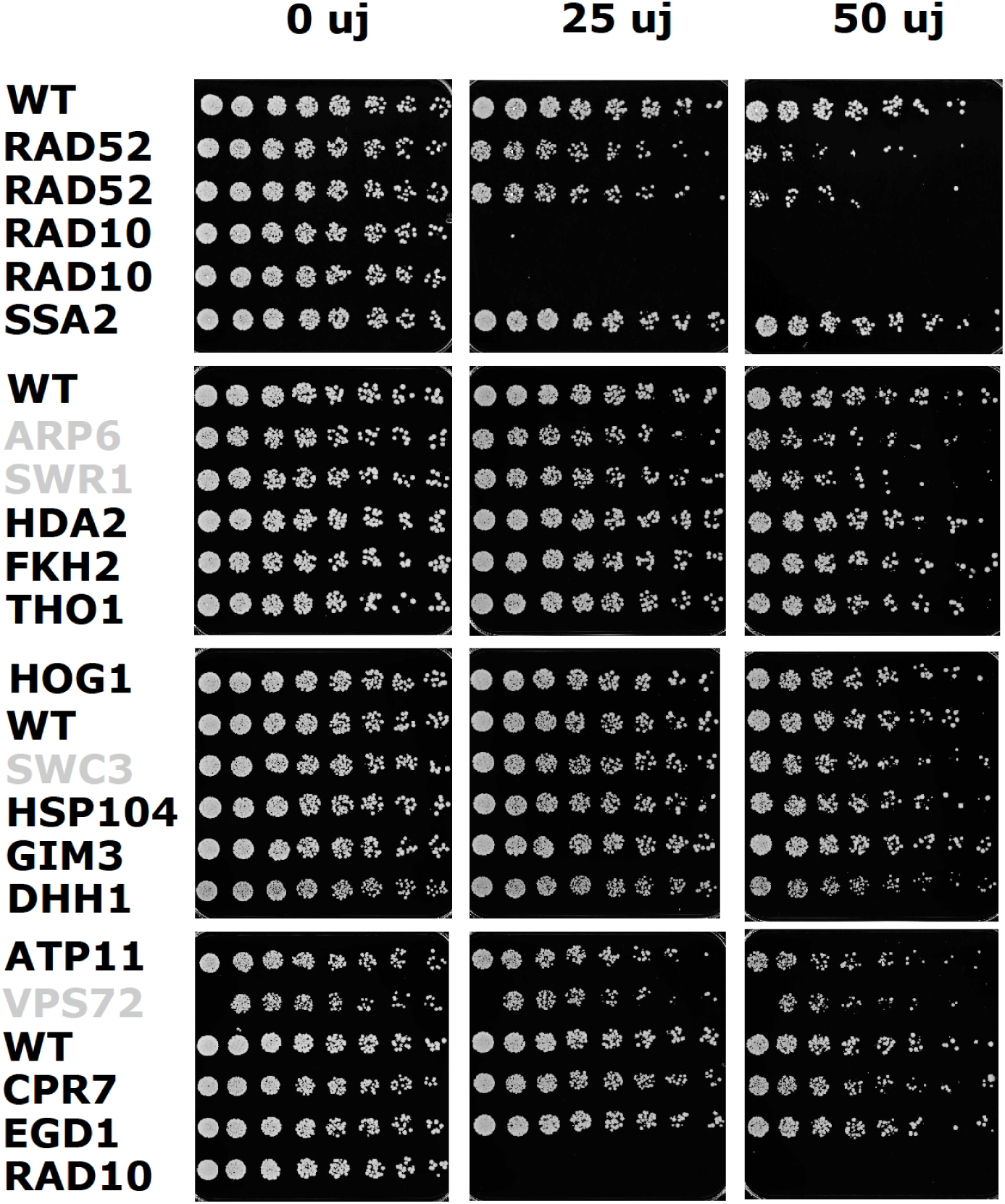
Spotting assay to test if deletions of candidate buffer genes increases UV sensitivity. Dilution series were spotted on YPD agar plates and exposed to 0, 25 or 50 uJ/cm^2^ of UV light. Each plate includes a WT strain, and positive controls *rad52* and *rad10* were included (which show clear increase in UV sensitivity). Deletion strains in grey were not selected as candidate genes, but are included to show unmodified pictures of full plates.

**Supplementary figure 3.**
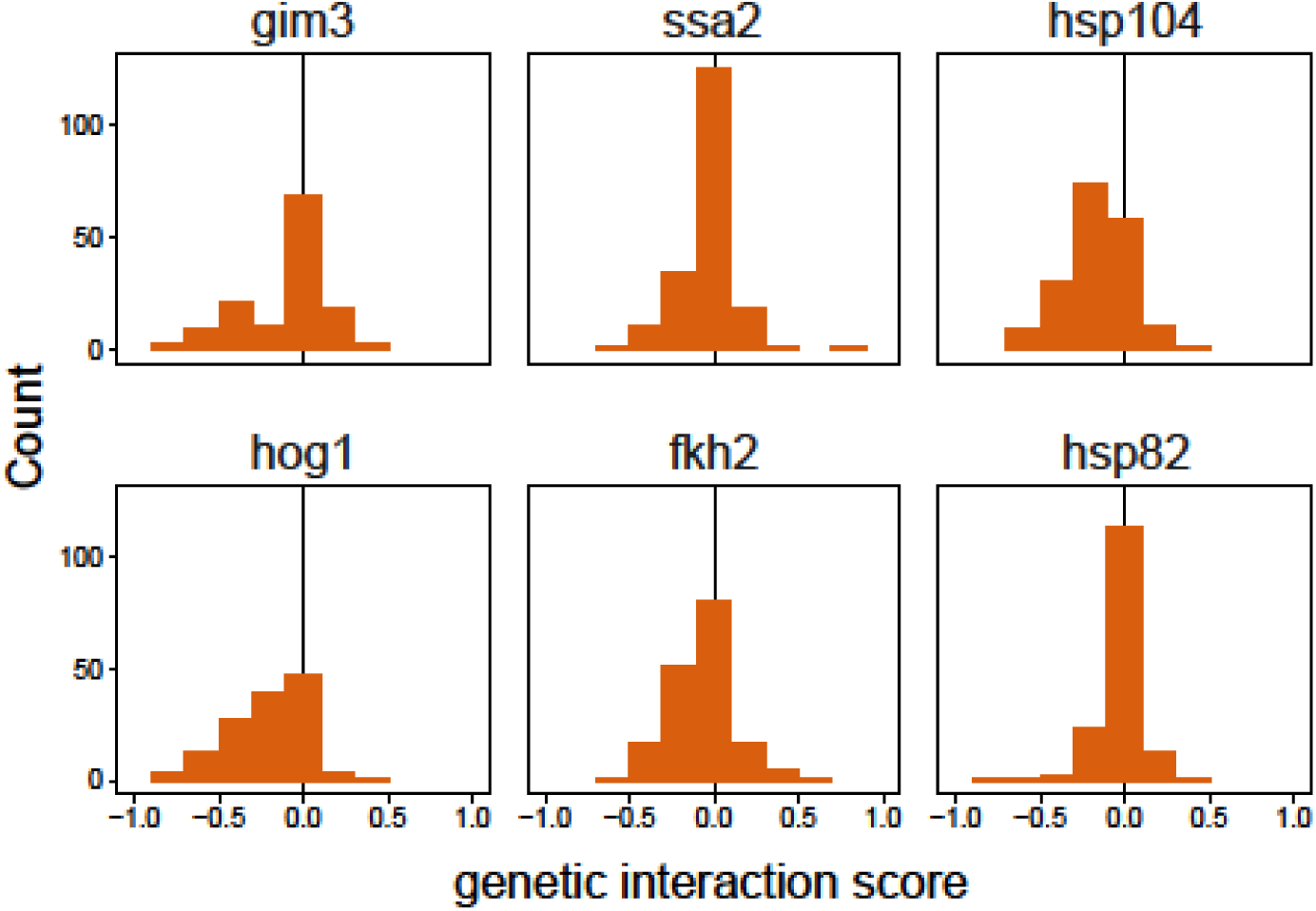
The genetic interaction scores of all tested mutant strains represented in Figure 4b split up by the tested buffer gene. Additionally, genetic interaction scores for *HSP82* is represented in this figure (which was excluded from Figure 4B).

**Supplementary figure 4.**
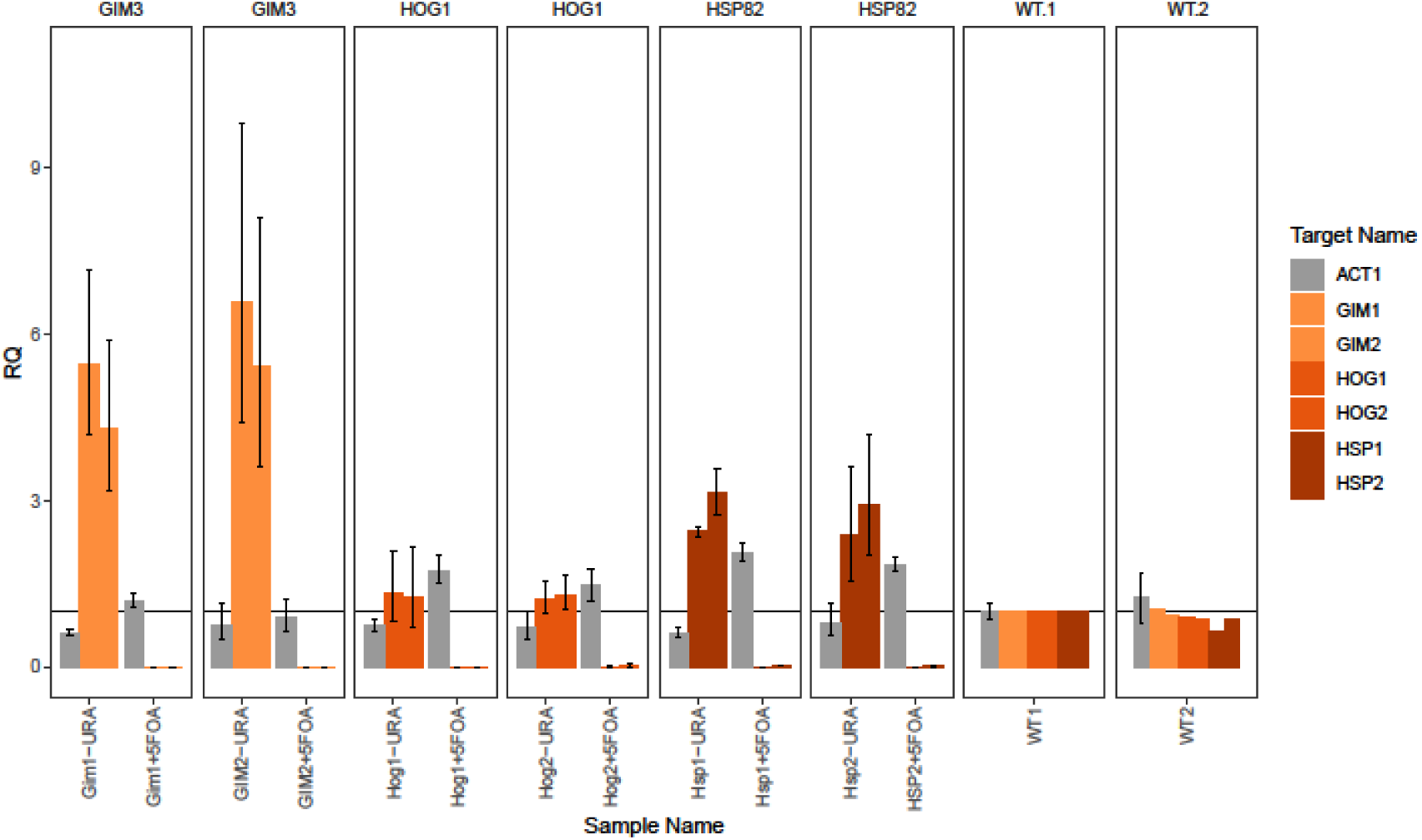
Gene expression determined by qPCR and relative to WT strain endogenous control gene *THD3.* Candidate genes (*GIM3*, *HOG1* and *HSP82*) were on a selectable and counter-selectable plasmid, and expression of the genes was measured after selecting for the plasmid (gene-URA, 2 technical replicates) or against the plasmid (gene+5FOA, 2 technical replicates). Additionally for each strain, we measured expression of *ACT1*. The two biological replicates for each candidate gene show no expression of the gene after selection against the plasmid. Error bars represent minimum and maximum relative quantity.

## REFERENCES

Aguilar-Rodríguez, J., Sabater-Muñoz, B., Montagud-Martínez, R., Berlanga, V., Alvarez-Ponce, D., Wagner, A., and Fares, M.A. (2016). The Molecular Chaperone DnaK Is a Source of Mutational Robustness. Genome Biol. Evol. 8, 2979–2991.

Barrett, R.D.H., and Schluter, D. (2008). Adaptation from standing genetic variation. Trends Ecol. Evol. 23, 38–44.

Berger, D., Bauerfeind, S.S., Blanckenhorn, W.U., and Schäfer, M.A. (2011). High temperatures reveal cryptic genetic variation in a polymorphic female sperm storage organ. Evolution 65, 2830–2842.

Bolger, A.M., Lohse, M., and Usadel, B. (2014). Trimmomatic: A flexible trimmer for Illumina sequence data. Bioinformatics 30, 2114–2120.

Broer, L., Ikram, M.A., Schuur, M., Destefano, A.L., Bis, J.C., Liu, F., Rivadeneira, F., Uitterlinden, A.G., Beiser, A.S., Longstreth, W.T., et al. (2011). Association of HSP70 and its co-chaperones with Alzheimer’s disease. J. Alzheimers. Dis. 25, 93–102.

Burga, A., Casanueva, M.O., and Lehner, B. (2011). Predicting mutation outcome from early stochastic variation in genetic interaction partners. Nature 480, 250–253.

Carpenter, A.E., Jones, T.R., Lamprecht, M.R., Clarke, C., Kang, I.H., Friman, O., Guertin, D.A., Chang, J.H., Lindquist, R.A., Moffat, J., et al. (2006). CellProfiler: Image analysis software for identifying and quantifying cell phenotypes. Genome Biol. 7, 1–11.

Casanueva, M.O., Burga, A., and Lehner, B. (2012). Fitness trade-offs and environmentally induced mutation buffering in isogenic C. elegans. Science 335, 82–85.

Chen, B., and Wagner, A. (2012). Hsp90 is important for fecundity, longevity, and buffering of cryptic deleterious variation in wild fly populations. BMC Evol. Biol. 12, 25.

Cingolani, P., Platts, A., Wang, L.L., Coon, M., Nguyen, T., Wang, L., Land, S.J., Lu, X., and Ruden, D.M. (2012). A program for annotating and predicting the effects of single nucleotide polymorphisms, SnpEff. Fly 6, 80–92.

Costanzo, M., VanderSluis, B., Koch, E.N., Baryshnikova, A., Pons, C., Tan, G., Wang, W., Usaj, M., Hanchard, J., Lee, S.D., et al. (2016). A global genetic interaction network maps a wiring diagram of cellular function. Science 353, aaf1420.

Cowen, L.E., and Lindquist, S. (2005). Hsp90 Potentiates the Rapid Evolution of New Traits: Drug Resistance in Diverse Fungi. Science 309, 2185–2189.

Dworkin, I., Palsson, A., Birdsall, K., and Gibson, G. (2003). Evidence that Egfr contributes to cryptic genetic variation for photoreceptor determination in natural populations of Drosophila melanogaster. Curr. Biol. 13, 1888–1893.

Fares, M.A., Ruiz-González, M.X., Moya, A., Elena, S.F., and Barrio, E. (2002). GroEL buffers against deleterious mutations. Nature 417, 398.

Gallone, B., Steensels, J., Prahl, T., Soriaga, L., Saels, V., Herrera-Malaver, B., Merlevede, A., Roncoroni, M., Voordeckers, K., Miraglia, L., et al. (2016). Domestication and Divergence of Saccharomyces cerevisiae Beer Yeasts. Cell 166, 1397–1410.

Geiler-Samerotte, K.A., Zhu, Y.O., Goulet, B.E., Hall, D.W., Siegal, M.L., Rutherford, S., Lindquist, S., Queitsch, C., Sangster, T., Lindquist, S., et al. (2016). Selection Transforms the Landscape of Genetic Variation Interacting with Hsp90. PLOS Biol. 14, e2000465.

Giaever, G., Chu, A.M., Ni, L., Connelly, C., Riles, L., Véronneau, S., Dow, S., Lucau-Danila, A., Anderson, K., André, B., et al. (2002). Functional profiling of the Saccharomyces cerevisiae genome. Nature 418, 387–391.

Gibson, G., and Dworkin, I. (2004a). Uncovering cryptic genetic variation. Nat. Rev. Genet. 5, 681–690.

Gibson, G., and Dworkin, I. (2004b). Uncovering cryptic genetic variation. Nat. Rev. Genet. 5, 681–690.

Gibson, G., and Hogness, D.S. (1996). Effect of Polymorphism in the Drosophila Regulatory Gene Ultrabithorax on Homeotic Stability. Science 271, 200–203.

Girstmair, H., Tippel, F., Lopez, A., Tych, K., Stein, F., Haberkant, P., Schmid, P.W.N., Helm, D., Rief, M., Sattler, M., et al. (2019). The Hsp90 isoforms from S. cerevisiae differ in structure, function and client range. Nat. Commun. 10, 3626.

Grote, S. (2021). GOfuncR: Gene ontology enrichment using FUNC. R package version 1.14.0.

Gruber, C., and Bogunovic, D. (2020). Incomplete penetrance in primary immunodeficiency: a skeleton in the closet. Hum. Genet. 139, 745–757.

Hayden, E.J., Ferrada, E., and Wagner, A. (2011). Cryptic genetic variation promotes rapid evolutionary adaptation in an RNA enzyme. Nature 474, 92–95.

Hietpas, R.T., Jensen, J.D., and Bolon, D.N.A. (2011). Experimental illumination of a fitness landscape. Proc. Natl. Acad. Sci. U. S. A. 108, 7896–7901.

Hummel, B., Hansen, E.C., Yoveva, A., Aprile-Garcia, F., Hussong, R., and Sawarkar, R. (2017). The evolutionary capacitor HSP90 buffers the regulatory effects of mammalian endogenous retroviruses. Nat. Struct. Mol. Biol. 24, 234–242.

Jarosz, D.F., and Lindquist, S. (2010). Hsp90 and Environmental Stress Transform the Adaptive Value of Natural Genetic Variation. Science 330, 1820–1824.

Karras, G.I., Yi, S., Sahni, N., Fischer, M., Xie, J., Vidal, M., D’Andrea, A.D., Whitesell, L., and Lindquist, S. (2017). HSP90 Shapes the Consequences of Human Genetic Variation. Cell 168, 856–866.e12.

Koneru, S.L., Hintze, M., Katsanos, D., and Barkoulas, M. (2021). Cryptic genetic variation in a heat shock protein modifies the outcome of a mutation affecting epidermal stem cell development in C. elegans. Nat. Commun. 12, 3263.

Lauter, N., and Doebley, J. (2002). Genetic Variation for Phenotypically Invariant Traits Detected in Teosinte: Implications for the Evolution of Novel Forms. Genetics 160, 333–342.

Lehner, B., Crombie, C., Tischler, J., Fortunato, A., and Fraser, A.G. (2006). Systematic mapping of genetic interactions in Caenorhabditis elegans identifies common modifiers of diverse signaling pathways. Nat. Genet. 38, 896–903.

Levy, S.F., Siegal, M.L., Wagner, A., Waddington, C., Siegal, M., Bergman, A., Hansen, T., Wagner, A., Queitsch, C., Sangster, T., et al. (2008). Network Hubs Buffer Environmental Variation in Saccharomyces cerevisiae. PLoS Biol. 6, e264.

Li, H., and Durbin, R. (2009). Fast and accurate short read alignment with Burrows–Wheeler transform. Bioinformatics 25, 1754–1760.

Liang, J., Xia, L., Oyang, L., Lin, J., Tan, S., Yi, P., Han, Y., Luo, X., Wang, H., Tang, L., et al. (2020). The functions and mechanisms of prefoldin complex and prefoldin-subunits. Cell Biosci. 10, 1–15.

Lurias, S.E., and Delbrock, M. (1943). Mutations of Bacteria from Virus Sensitivity to Virus Resistance. Genetics 28, 491–511.

Maisnier-Patin, S., Roth, J.R., Fredriksson, Å., Nyström, T., Berg, O.G., and Andersson, D.I. (2005). Genomic buffering mitigates the effects of deleterious mutations in bacteria. Nat. Genet. 37, 1376–1379.

March, Z.M., Sweeney, K., Kim, H., Yan, X., Castellano, L.M., Jackrel, M.E., Lin, J., Chuang, E., Gomes, E., Willicott, C.W., et al. (2020). Therapeutic genetic variation revealed in diverse Hsp104 homologs. Elife 9, 1–52.

McGuigan, K., Nishimura, N., Currey, M., Hurwit, D., and Cresko, W.A. (2011). Cryptic genetic variation and body size evolution in threespine stickleback. Evolution 65, 1203–1211.

McKenna, A., Hanna, M., Banks, E., Sivachenko, A., Cibulskis, K., Kernytsky, A., Garimella, K., Altshuler, D., Gabriel, S., Daly, M., et al. (2010). The Genome Analysis Toolkit: a MapReduce framework for analyzing next-generation DNA sequencing data. Genome Res. 20, 1297–1303.

Millson, S.H., Truman, A.W., King, V., Prodromou, C., Pearl, L.H., and Piper, P.W. (2005). A two-hybrid screen of the yeast proteome for Hsp90 interactors uncovers a novel Hsp90 chaperone requirement in the activity of a stress-activated mitogen-activated protein kinase, Slt2p (Mpk1p). Eukaryot. Cell 4, 849–860.

Paaby, A.B., and Rockman, M. V. (2014). Cryptic genetic variation: evolution’s hidden substrate. Nat. Rev. Genet. 15, 247–258.

Queitsch, C., Sangster, T.A., and Lindquist, S. (2002). Hsp90 as a capacitor of phenotypic variation. Nature 417, 618–624.

R Core Team (2021). R Core Team. R A Lang. Environ. Stat. Comput.

Richardson, J.B., Uppendahl, L.D., Traficante, M.K., Levy, S.F., Siegal, M.L., Masel, J., Siegal, M., Gibson, G., Whitesell, L., Lindquist, S., et al. (2013). Histone Variant HTZ1 Shows Extensive Epistasis with, but Does Not Increase Robustness to, New Mutations. PLoS Genet. 9, e1003733.

Rohner, N., Jarosz, D.F., Kowalko, J.E., Yoshizawa, M., Jeffery, W.R., Borowsky, R.L., Lindquist, S., and Tabin, C.J. (2013). Cryptic Variation in Morphological Evolution: HSP90 as a Capacitor for Loss of Eyes in Cavefish. Science 342, 1372–1375.

Rstudio, T. (2021). RStudio: Integrated Development for R. Rstudio Team, PBC, Boston, MA.

Rutherford, S.L. (2003). Between genotype and phenotype: protein chaperones and evolvability. Nat. Rev. Genet. 4, 263–274.

Rutherford, S.L., and Lindquist, S. (1998). Hsp90 as a capacitor for morphological evolution. Nature 396, 336–342.

Sangster, T.A., Salathia, N., Undurraga, S., Milo, R., Schellenberg, K., Lindquist, S., and Queitsch, C. (2008). HSP90 affects the expression of genetic variation and developmental stability in quantitative traits. Proc. Natl. Acad. Sci. USA 105, 2963–2968.

Siegal, M.L. (2017). Molecular genetics: Chaperone protein gets personal. Nature 545, 36–37.

Siegal, M.L., and Leu, J.-Y. (2014). On the Nature and Evolutionary Impact of Phenotypic Robustness Mechanisms. Annu. Rev. Ecol. Evol. Syst. 45, 495–517.

Stark, C., Breitkreutz, B.J., Reguly, T., Boucher, L., Breitkreutz, A., and Tyers, M. (2006). BioGRID: a general repository for interaction datasets. Nucleic Acids Res. 34, D535–D539.

Tahmaz, I., Shahmoradi Ghahe, S., and Topf, U. (2022). Prefoldin Function in Cellular Protein Homeostasis and Human Diseases. Front. Cell Dev. Biol. 9, 3949.

Tirosh, I., Reikhav, S., Sigal, N., Assia, Y., and Barkai, N. (2010). Chromatin regulators as capacitors of interspecies variations in gene expression. Mol. Syst. Biol. 6, 435.

Tokuriki, N., and Tawfik, D.S. (2009). Chaperonin overexpression promotes genetic variation and enzyme evolution. Nature 459, 668–673.

Tong, A.H.Y., Lesage, G., Bader, G.D., Ding, H., Xu, H., Xin, X., Young, J., Berriz, G.F., Brost, R.L., Chang, M., et al. (2004). Global Mapping of the Yeast Genetic Interaction Network. Science 303, 808–813.

Valiadi, M., Debora Iglesias-Rodriguez, M., and Amorim, A. (2012). Distribution and genetic diversity of the luciferase gene within marine dinoflagellates. J. Phycol. 48, 826–836.

Voordeckers, K., Kominek, J., Das, A., Espinosa-Cantú, A., De Maeyer, D., Arslan, A., Van Pee, M., van der Zande, E., Meert, W., Yang, Y., et al. (2015). Adaptation to High Ethanol Reveals Complex Evolutionary Pathways. PLOS Genet. 11, e1005635.

Voordeckers, K., Colding, C., Grasso, L., Pardo, B., Hoes, L., Kominek, J., Gielens, K., Dekoster, K., Gordon, J., Van der Zande, E., et al. (2020). Ethanol exposure increases mutation rate through error-prone polymerases. Nat. Commun. 11, 3664.

Vyas, V.K., Bushkin, G.G., Bernstein, D.A., Getz, M.A., Sewastianik, M., Barrasa, M.I., Bartel, D.P., and Fink, G.R. (2018). New CRISPR Mutagenesis Strategies Reveal Variation in Repair Mechanisms among Fungi. mSphere 3, e00154–18.

Waddington, C. (1942). Canalization of development and the inheritance of acquired characters. Nature 150, 563–565.

Waddington, C.H. (1953). Genetic assimialtion of an acquired character. Evolution 7, 118–126.

Wagih, O., and Parts, L. (2014). gitter: a robust and accurate method for quantification of colony sizes from plate images. G3 *4*, 547–552.

Wagih, O., Usaj, M., Baryshnikova, A., VanderSluis, B., Kuzmin, E., Costanzo, M., Myers, C.L., Andrews, B.J., Boone, C.M., and Parts, L. (2013). SGAtools: one-stop analysis and visualization of array-based genetic interaction screens. Nucleic Acids Res. 41, W591–W596.

Westfall, P.J., Ballon, D.R., and Thorner, J. (2004). When the stress of your environment makes you go HOG wild. Science 306, 1511–1512.

Winzeler, E.A., Shoemaker, D.D., Astromoff, A., Liang, H., Anderson, K., Andre, B., Bangham, R., Benito, R., Boeke, J.D., Bussey, H., et al. (1999). Functional characterization of the S. cerevisiae genome by gene deletion and parallel analysis. Science 285, 901–906.

Yang, X.X., Maurer, K.C., Molanus, M., Mager, W.H., Siderius, M., and Van Der Vies, S.M. (2006). The molecular chaperone Hsp90 is required for high osmotic stress response in Saccharomyces cerevisiae. FEMS Yeast Res. 6, 195–204.

Yeyati, P.L., Bancewicz, R.M., Maule, J., and Van Heyningen, V. (2007). Hsp90 Selectively Modulates Phenotype in Vertebrate Development. PLOS Genet. 3, e43.

Zabinsky, R.A., Mason, G.A., Queitsch, C., and Jarosz, D.F. (2019). It’s not magic – Hsp90 and its effects on genetic and epigenetic variation. Semin. Cell Dev. Biol. 88, 21–35.

Zhao, R., Davey, M., Hsu, Y.C., Kaplanek, P., Tong, A., Parsons, A.B., Krogan, N., Cagney, G., Mai, D., Greenblatt, J., et al. (2005). Navigating the chaperone network: An integrative map of physical and genetic interactions mediated by the hsp90 chaperone. Cell 120, 715–727.

Zheng, Q. (2017). rSalvador: An R Package for the Fluctuation Experiment. G3 *7*, 3849–3856.

